# Sex-specific differences in the function and differentiation of ABCs mark TLR7-driven immunopathogenesis

**DOI:** 10.1101/2021.01.20.427400

**Authors:** Edd Ricker, Michela Manni, Danny Flores-Castro, Daniel Jenkins, Sanjay Gupta, Juan Rivera-Correa, Wenzhao Meng, Aaron M. Rosenfeld, Tania Pannellini, Mahesh Bachu, Yurii Chinenov, Peter K. Sculco, Rolf Jessberger, Eline T. Luning Prak, Alessandra B. Pernis

## Abstract

Sex differences characterize immune responses to viruses and autoimmune diseases like SLE. ABCs are an emerging population of CD11c^+^ T-bet^+^ B cells critical for antiviral responses and autoimmune disorders. DEF6 and SWAP70, are two homologous molecules whose combined absence in double-knock-out mice (DKOs) leads to a lupus syndrome in females marked by an accumulation of ABCs. Here we demonstrate that DKO ABCs exhibit sex-specific differences in their expansion, upregulation of an ISG signature, and further differentiation. BCR sequencing and fate mapping experiments reveal that DKO ABCs undergo oligoclonal expansion and differentiate into both CD11c^+^ and CD11c^-^ effector populations with pathogenic and proinflammatory potential. *Tlr7* duplication in DKO males overrides the sex-bias and further augments the dissemination and pathogenicity of ABCs resulting in severe pulmonary inflammation and early mortality. Thus, sexual dimorphism shapes the expansion, function, and differentiation of ABCs contributing to the sex-bias that accompanies TLR7-driven immunopathogenesis.

## INTRODUCTION

Sex-dependent differences in immune responses have been well documented in viral infections, vaccination outcomes, and autoimmune diseases like Systemic Lupus Erythematosus (SLE), a heterogeneous disorder that often includes upregulation of interferon stimulated genes (ISGs) in addition to autoantibody production and multi-organ involvement^1, 2^. Both sex hormones and genes on the X-chromosome have been implicated in this sexual dimorphism^3^. Notably, TLR7, an endosomal TLR critical for responses to viruses like SARS-CoV-2 and lupus pathogenesis, is encoded on the X-chromosome and has recently been shown to partially escape X-chromosome inactivation resulting in greater TLR7 expression in a proportion of female B cells, monocytes, and pDCs^4^.

TLR7 engagement promotes the formation of Age/autoimmune-associated B cells (ABCs), also known as DN2 in humans, a B cell population that preferentially expands with age in female mice^5,6,7^. ABCs exhibit a distinctive phenotype and, in addition to classical B cell markers, express the transcription factor T-bet and myeloid markers like CD11c. T-bet and CD11c are often, but not always, co-expressed^8,9,10^. ABCs are an important component of antiviral responses and are inappropriately controlled in several viral infections including HIV and SARS-Co-V2^11,12,13^. Aberrant expansion and activation of ABCs is also associated with autoimmune pathogenesis, especially SLE^14^. In this disease, ABCs accumulate to a greater extent in African-American patients, rapidly differentiate into plasmablasts/plasma cells (PB/PC), are major producers of autoantibodies, and correlate with disease activity and clinical manifestations^7, 15, 16^. Despite the emerging biological and clinical importance of ABCs, the full spectrum of their function and differentiation capabilities are incompletely understood.

Although T-bet is a well-known marker for ABCs, reliance of ABCs on this transcription factor differs depending on the setting. B-cell T-bet is important for protective flu-specific IgG2c antibodies, but its absence has variably impacted the generation of ABCs and disease parameters in lupus murine models^9, 17,18,19^. This is likely due to the presence of additional regulators of autoimmune ABCs such as IRF5 (Interferon Regulatory Factor 5), whose dysregulation promotes ABC accumulation and lupus development^20^. The ability of IRF5 to drive ABC expansion can be restrained by the SWEF proteins, Def6 and SWAP-70, two homologous proteins that also control cytoskeletal reorganization by regulating Rho GTPase signaling and whose combined absence in mice results in a lupus syndrome that primarily affects females^21,22,24^. An important role for these molecules in immune responses, inflammation, and autoimmunity is supported not only by murine genetic models but also by human studies. The CORO1A-DEF6 blood transcription module correlates with responses to flu vaccination and malaria^25, 26^. Furthermore, *SWAP70* is a susceptibility locus for RA^27^ and CVD^28^ while *DEF6* is a risk factor for human SLE^29, 30^. Mutations in *DEF6* moreover result in early-onset autoimmune manifestations, often associated with viral infections, which include autoantibody production and upregulation of an ISG signature^31, 32^.

In this study we have exploited the sex-bias exhibited by mice lacking both SWEF proteins (Double-Knock-out or DKOs) to investigate the impact of sexual dimorphism on the ABC compartment. We demonstrate that ABCs from DKO females and males differ in their ability to expand, upregulate an ISG signature, and further differentiate. BCR sequencing and fate mapping experiments reveal marked oligoclonal expansion and interrelatedness of ABCs with both CD11c^+^ and CD11c^-^ effector populations, which include CD11c^+^ pre-GC B cells and CD11c^+^ PBs. In addition to IRF5, DKO ABCs also require IRF8 but are less dependent on T-bet.

Notably, *Tlr7* duplication in DKO males overrides the sex-bias and augments the pathogenicity of ABCs resulting in severe pathology and early mortality. Thus, in autoimmune settings, ABCs can give rise to a heterogenous population of effector cells with distinct pathogenic potentials that are controlled in a sexually dimorphic manner.

## RESULTS

### ABC accumulation and function in DKOs is sex-dependent and controlled by TLR7

Similar to human SLE, the lupus syndrome that develops in DKOs preferentially affects females providing a powerful model to delineate the cellular and molecular mechanisms that underlie sexual dimorphism in autoimmunity. Given the key role of ABCs in lupus, we first assessed whether the sex-bias that accompanies lupus development in DKOs was associated with differences in ABC expansion. Significantly more ABCs accumulated in DKO females than age-matched DKO males, although DKO males still contained greater numbers of ABCs than WT controls (Fig. 1A). Furthermore, ABCs sorted from DKO males secreted significantly lower levels of anti-dsDNA IgG2c upon stimulation with a TLR7 agonist, imiquimod, than ABCs from DKO females (Fig. 1B). Thus, both the accumulation and the function of ABCs in DKOs are controlled in a sex-specific manner.

**Figure 1.**
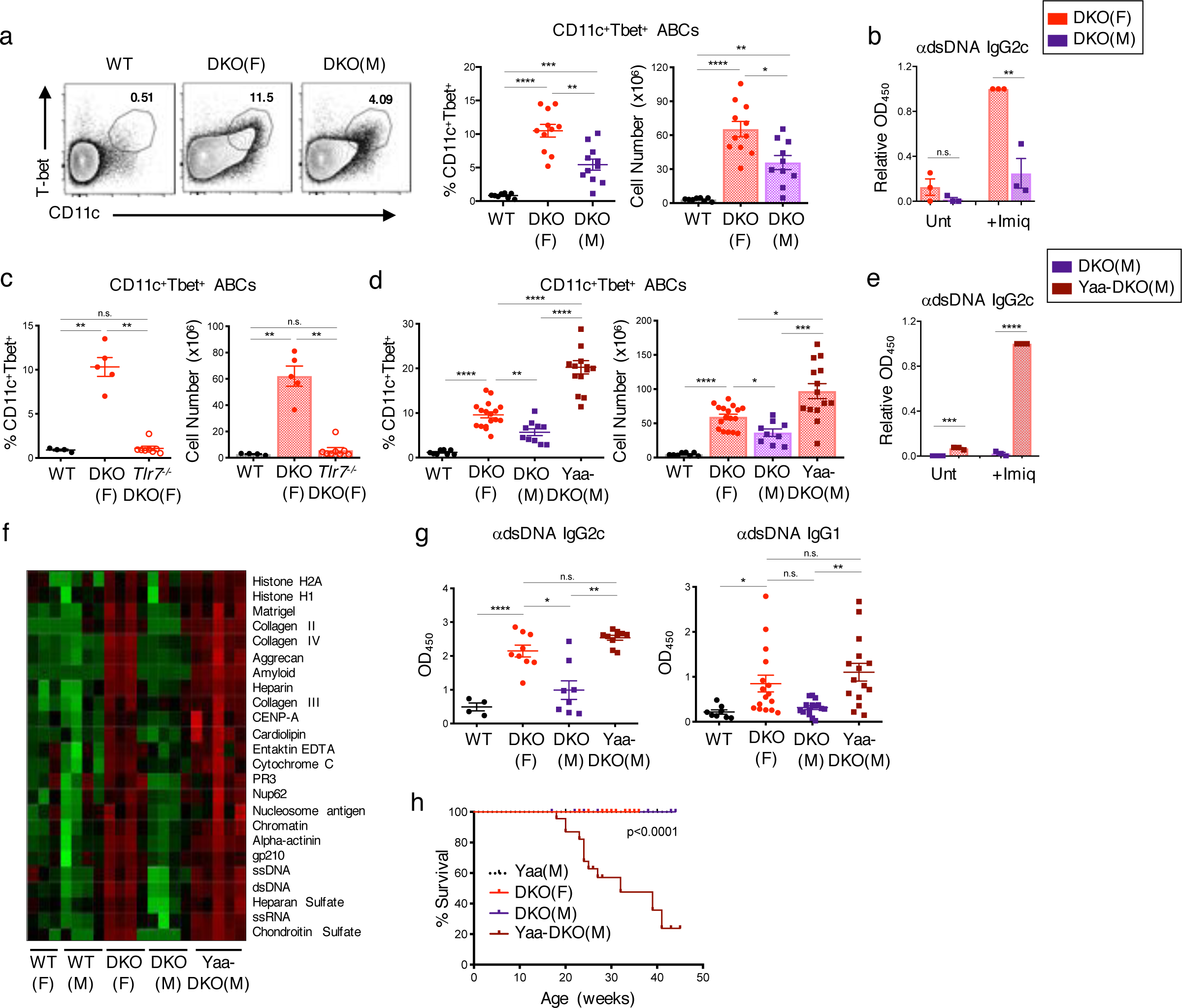
Sex-specific differences in ABC accumulation and function in DKO mice are controlled by TLR7. (A) Representative FACS plots and quantifications of CD11c^+^Tbet^+^ ABCs (gated on B220^+^) from the spleens of aged (20-30wk) WT, female DKO (F), and male DKO (M) mice. Data shows mean +/-SEM; *n*>=8 per genotype; p-value by Brown-Forsythe and Welch ANOVA followed by Games-Howell’s test for multiple comparisons. (B) Pooled ELISA data for anti-dsDNA IgG2c antibodies from supernatants of sorted ABCs (B220^+^CD19^+^CD11c^+^CD11b^+^) from DKO(F) or DKO(M) mice after culturing with Imiquimod for 7d. Data shows mean +/-SEM; *n*=3; p-value by unpaired two-tailed t-tests. (C) Quantifications of CD11c^+^Tbet^+^ ABCs from the spleens of WT, DKO(F), and *Tlr7^-/-^.*DKO(F) mice. Data show mean +/-SEM; *n*>=4 per genotype; p-value by Brown-Forsythe and Welch ANOVA followed by Games-Howell’s test for multiple comparisons. (D) Quantifications of CD11c^+^Tbet^+^ ABCs (gated on B220^+^) from the spleens of aged (24-30wk) WT, DKO(F), DKO(M), and Yaa-DKO(M) mice. Data shows mean +/- SEM; *n*>=8 per genotype; p-value by Brown-Forsythe and Welch ANOVA followed by Games-Howell’s test for multiple comparisons. (E) Pooled ELISA data for anti-dsDNA IgG2c antibodies from supernatants of sorted ABCs from DKO(M) or Yaa-DKO(M) mice after culturing with Imiquimod for 7d. Data shows mean +/- SEM; *n*=3; p-value by unpaired two-tailed t-tests. (F) Autoantigen microarray showing the relative autoantibody levels in the serum of female WT (F), male WT (M), DKO(F), DKO(M), and Yaa-DKO(M) mice. Data shows serum from at least 3 mice per genotype. (G) Pooled ELISA data for anti-dsDNA IgG2c and IgG1 in the serum from the indicated mice. Data shows mean +/- SEM; *n*>=4 per genotype; p-value by Brown-Forsythe and Welch ANOVA followed by Games-Howell’s test for multiple comparisons. (H) Plot showing survival rate of Yaa control *(black)*, DKO(F) *(red)*, DKO(M) *(purple)*, and Yaa-DKO(M) *(maroon)* mice across 45 weeks. Data represents a cohort of 14 Yaa control, 16 DKO(F), 9 DKO(M), and 31 Yaa-DKO(M) mice; p-value by Mantel-Cox test. *p <0.05, **p <0.01, ***p <0.001, ****p <0.0001.

*Tlr7* can be expressed biallelically in a proportion of female B cells due to incomplete X chromosome inactivation^4^. In line with these findings, ABCs from DKO females expressed higher levels of *Tlr7* than ABCs from DKO males (Fig. S1A). ABC accumulation in DKO females was furthermore dependent on TLR7, as DKO females crossed to *Tlr7*^-/-^ mice exhibited a profound reduction in ABC accumulation (Fig. 1C). *Tlr7*-deficient DKO females also displayed significant decreases in GC B cells, PB/PCs, and TFH cells, and lacked anti-dsDNA IgG2c antibodies (Fig. S1B-E). Thus, ABC expansion and lupus pathogenesis in DKO females are dependent on *Tlr7*.

To further assess the importance of *Tlr7* in the sex-bias of DKOs, we crossed DKO males to C57BL/6 mice carrying the Y-linked genomic modifier *Yaa* (termed Yaa-DKOs), in which a portion of the X-chromosome has translocated onto the Y-chromosome resulting in a 2-fold increase in Tlr7 expression in males^33^. *Tlr7* duplication in DKO males markedly increased the frequencies and numbers of splenic ABCs reaching levels that were even greater than those observed in DKO females (Fig. 1D; S1F). *Tlr7* duplication in DKO males also rescued the ability of sorted male ABCs to secrete anti-dsDNA IgG2c antibodies upon stimulation (Fig. 1E). Increased ABC accumulation and function in Yaa-DKO males were accompanied by autoantibody production, the classical clinical feature of SLE (Fig. 1F-G). Total antibody titers were also comparable between DKO females and Yaa-DKO males (Fig. S1G). Yaa-DKO males also exhibited significantly decreased survival as compared to both DKO males and females (Fig. 1H). Thus, duplication of *Tlr7* in Yaa-DKO males overrides the sex-bias and promotes the development of a severe lupus syndrome in DKO males marked by greatly enhanced accumulation of ABCs and autoantibody responses.

### The expansion of GC B cells and PB/PCs in DKOs is regulated in a sex-specific manner

In addition to ABC accumulation, DKO females also exhibit robust GC and PB/PC responses^22^, prompting us to examine whether sex-specific differences could also be observed in these compartments. DKO females contained more GL7^+^Fas^+^ GC B cells than DKO males, a difference that was again reversed by *Tlr7* duplication in Yaa-DKO males (Fig. 2A). Immunofluorescence staining confirmed these findings and revealed that GCs in Yaa-DKO males were smaller and less well-organized than those in DKO females (Fig. 2B; S2A). DKO females also demonstrated a greater expansion of PB/PCs than DKO males (Fig. 2C). *Tlr7* duplication in Yaa-DKO males reversed this effect and strongly promoted the accumulation of PB/PCs, which was primarily observed in the spleen but not in the BM (Fig. 2C; S2B). No sex-based differences were detected in other B cell compartments (Fig. S2C-E). Other parameters known to promote spontaneous GC responses such as the ratio between TFH and TFR or the dual production of IFN*γ* and IL-21 were comparable between DKO females and males and only minimally affected by *Tlr7* duplication (Fig. 2D-E; S2F-G). Thus, DKOs exhibit a sex-specific accumulation of GC B cells and PB/PCs, which can be regulated in a *Tlr7-*dependent manner.

**Figure 2.**
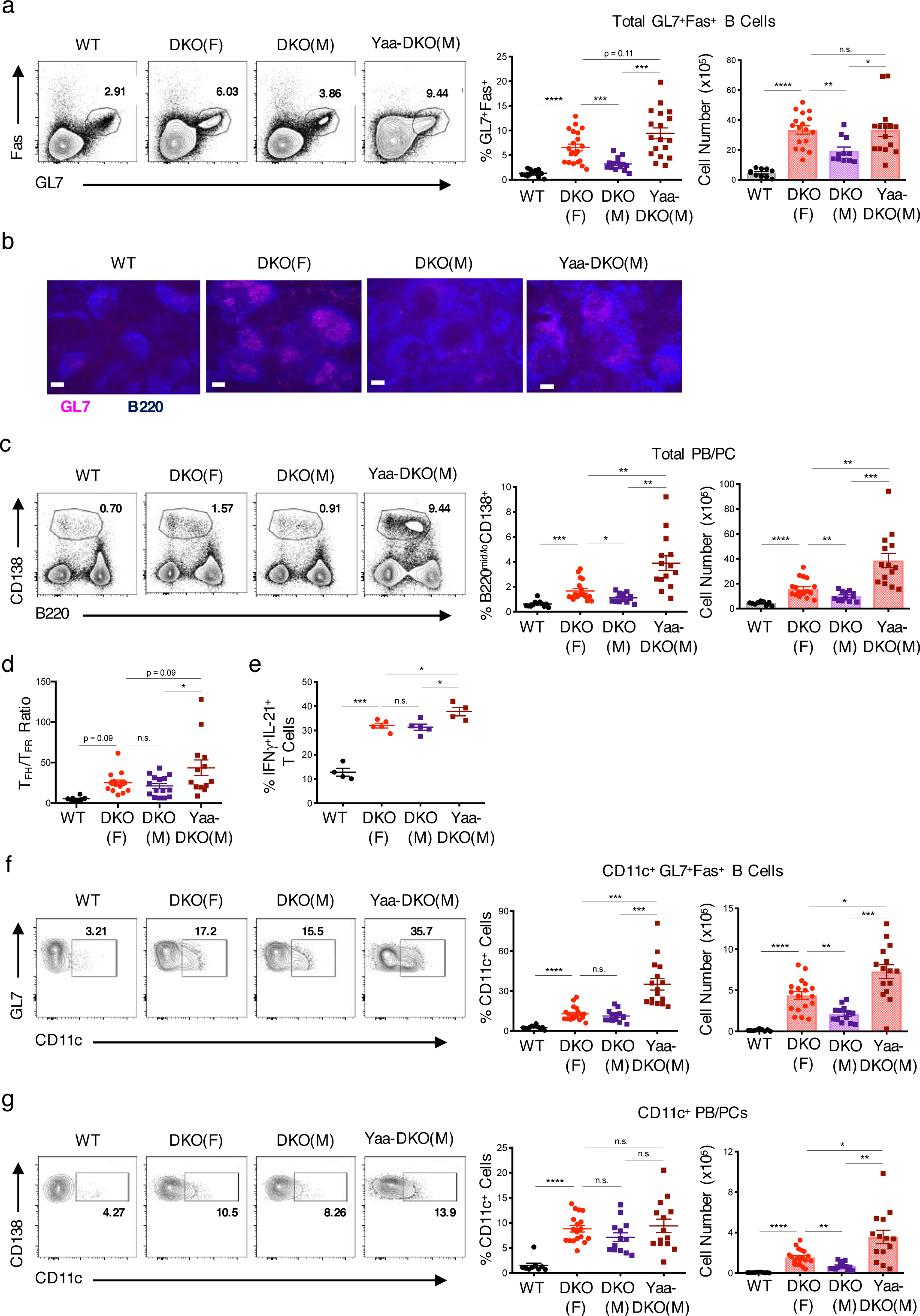
Sex-dependent accumulation of GC B cells and PB/PCs in DKO mice. (A) Representative FACS plots and quantifications of total GC B cells (gated on B220^+^ GL7^+^Fas^+^ splenocytes) from WT, female DKO(F), male DKO(M), and Yaa-DKO(M) mice. Data show mean +/- SEM; *n*>=14 per genotype; p-value by Brown-Forsythe and Welch ANOVA followed by Games-Howell’s test for multiple comparisons. (B) Representative immunofluorescence images of B220^+^ (*blue)* GL7^+^ *(pink)* GCs in spleens from the indicated mice. Data representative of at least 4 frames for at least 2 mice per genotype. Bars show 50 μm. (C) Representative FACS plots and quantifications of total PB/PC (B220^mid/lo^CD138^+^) in spleens of the indicated mice. Data show mean +/- SEM; *n*>=13 per genotype; p-value by Brown-Forsythe and Welch ANOVA followed by Games-Howell’s test for multiple comparisons. (D) Quantification of the FACS plots showing T_FH_ (CD4^+^ PD1^hi^CXCR5^+^ Foxp3^-^) to T_FR_ (CD4^+^ PD1^hi^CXCR5^+^ Foxp3^+^) ratio in spleens from the indicated mice. Data show mean +/- SEM; *n*>=8 per genotype; p-value by Brown-Forsythe and Welch ANOVA followed by Games-Howell’s test for multiple comparisons. (E) Quantification of FACS plots showing IFN*γ*^+^IL-21^+^ T cells in spleens of the indicated mice following 4hr treatment of splenocytes with PMA/Ionomycin. Data show mean +/- SEM; *n*>=4 per genotype; p-value by Brown-Forsythe and Welch ANOVA followed by Games-Howell’s test for multiple comparisons. (F) Representative FACS plots and quantifications of CD11c^+^ GL7^+^Fas^+^ cells (gated on B220^+^ splenocytes) from the indicated mice. Data show mean +/- SEM; *n*>=11 per genotype; p-value by Brown-Forsythe and Welch ANOVA followed by Games- Howell’s test for multiple comparisons. (G) Representative FACS plots and quantifications of CD11c^+^ PB/PC (B220^mid/lo^CD138^+^) from the indicated mice. Data show mean +/- SEM; *n*>=9 per genotype; p-value by Brown-Forsythe and Welch ANOVA followed by Games-Howell’s test for multiple comparisons. *p <0.05, **p <0.01, ***p <0.001, ****p <0.0001.

Given the sex-bias in the accumulation of ABCs as well as of GC B cells and PB/PCs, we next investigated whether these populations might be related. An analysis of the GC B cell population in DKO females revealed that a subset of these cells expressed CD11c and that the numbers of CD11c^+^ GC B cells were significantly greater in DKO females than DKO males or WT controls (Fig. 2F; S2H). CD11c^+^ GC B cells were greatly increased in Yaa-DKO males (Fig. 2F). We also identified a population of CD11c^+^ PB/PCs that accumulated in DKO females and, to an even greater extent, in Yaa-DKO males (Fig. 2G; S2I). Thus, sex differences in lupus development in DKOs are accompanied by the aberrant accumulation of CD11c-expressing B cell effector subsets.

### Oligoclonal expansion and interrelatedness of ABCs and CD11c^+^ and CD11c^-^ B cell effector populations

To gain further insights into the effector B cell subsets that differentially expand in the spleens of DKO mice, we next compared their BCR repertoires. To evaluate the clonal landscape, we began by determining the contribution of the top 20 ranked clones to the overall repertoire by computing the D20 index^34^. We observed that the D20 index, or fraction of sequence copies contributing to the sum of the top 20 ranked clones, was lowest in FoBs and increased in ABCs, followed by GCB and finally being highest (most expanded) in the PB/PC pool (Fig. 3A). This order of large clone contribution by B cell subset was preserved in both DKO females and Yaa-DKO males (Fig. S3A). Furthermore, when one studies the level of resampling of clones between replicate sequencing libraries as an independent measure of clone size, the same trend is preserved, with FoBs having the lowest degree of overlap and PB/PCs having the highest (Fig. 3B; S3B). We next analyzed the level of somatic hypermutation (SHM), which revealed that GCB and PB/PC fractions had the highest frequencies of clones with SHM, while the ABCs had a level of SHM that was intermediate between FoBs and GCB/PBs (Fig. 3C; S3C-D). The SHM distribution trended by B cell subset rather than by mouse strain and the relative levels of SHM were preserved across these different B cell subsets irrespective of whether clones were unweighted or weighted by size (Fig. 3D; S3C-D). Given the somewhat lower levels of SHM in the ABCs as compared to the other B cell populations, we next turned to the length of the third complementary determining region (CDR3) and heavy chain variable (VH) gene usage as other general repertoire features. This analysis revealed longer CDR3 lengths for FoBs compared to the other B cell populations and VH gene usage that differed between FoBs and the other subsets (Fig. 3E-F). Taken together, this initial global repertoire analysis revealed that general repertoire features tended to map by B cell subset rather than by mouse strain, with FoBs having the greatest diversity, smallest clone size, and lowest level of SHM and GCB/PBs having the highest. ABCs instead tended to be intermediate with most of these measures.

**Figure 3.**
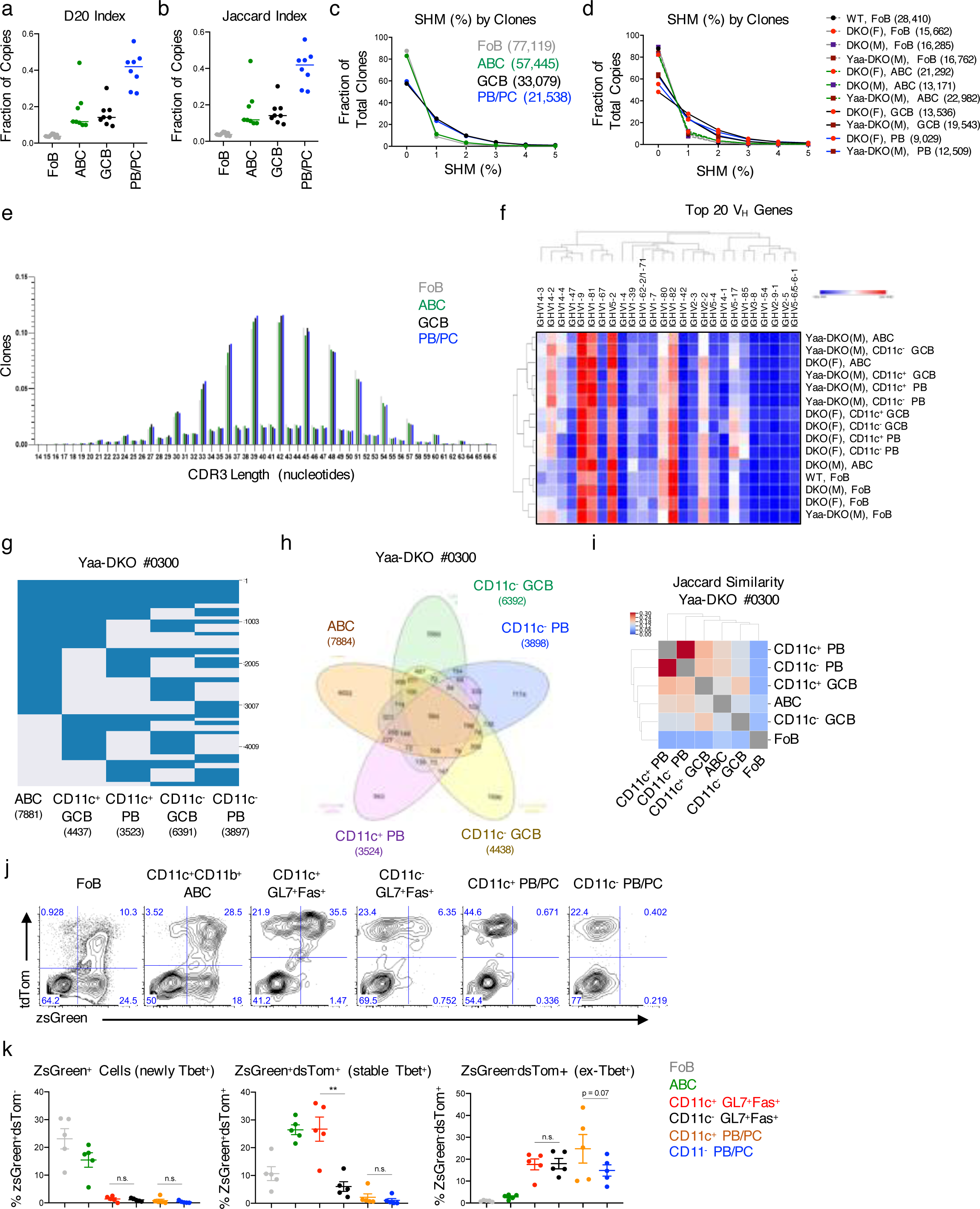
ABCs and CD11c^+^ and CD11c^-^ B cell effector populations in DKOs exhibit a high degree of clonal expansion and interrelatedness. (A) D20 index of each subset. D20 indicates the number of total copies in the top 20 clones as a fraction of total copies across all clones. Each dot represents a mouse. (B) Plot showing the level of overlap between clones in different sequencing libraries prepared from the same sample. For each comparison, each clone was only counted once (no weighting for clone size) and functional overlap was computed using the Jaccard Index. Each dot represents a mouse/subset sample. (C) Plot showing somatic hypermutation (SHM) as a fraction of total clones for each subset aggregated across mouse strains. The numbers in parenthesis indicate the total number of clones in the given subset. Each clone counts once for each subset. If a clone overlaps in multiple subsets, the SHM for each subset was calculated just for the sequences in a given subset. (D) The SHM by clones is aggregated across each mouse strain/subset combination as in Fig. 3C. (E) Plot showing CDR3 length (in nucleotides) of clones aggregated across all strains. Each clone is counted once in each subset/strain combination. (F) Heatmap showing usage of the 20 most frequent VH genes. Each clone is counted only once. Data are normalized by row and visualized in Morpheus using the default settings. (G) Plot showing clones (rows) that overlap between at least two B cell subsets (columns). Numbers along the right side of the plot indicate clone counts. Data from a single analysis of subsets from a Yaa-DKO mouse. (H) Venn diagram showing clonal overlap where numbers indicate clone counts in the different subset interactions. Data from a single analysis of subsets from a Yaa-DKO mouse. (I) Heatmap showing Jaccard similarity (fraction of clones that overlap between different two-subset comparisons). Diagonal values are excluded for scaling. Data from a single analysis of subsets from a Yaa-DKO mouse. (J-K) Female ZTCE-DKO mice were treated with tamoxifen to mark Tbet-expressing cells with tdTomato expression for 3d. Representative FACS plots *(J)* and quantifications *(K)* of zsGreen and tdTomato expression from the indicated populations. Data representative of and/or pooled from 5 mice and show mean +/- SEM; p-value by unpaired two-tailed t-tests. *p <0.05, **p <0.01, ***p <0.001, ****p <0.0001.

To further assess the relationships of ABCs with the other B cell effector populations that aberrantly expand in DKO females and Yaa-DKO males, we sorted and analyzed the various populations from individual mice, stratifying the GCB and PB/PC subsets by CD11c expression. CD11c is differentially expressed on DN B cells in human lupus and we wondered if similar interconnections existed in the context of the DKO females and Yaa-DKO male models. In particular, CD11c^+^ DN2 cells are transcriptionally and epigenetically poised to become PBs in human SLE^35^. If this also occurs in these models of murine SLE, then CD11c^+^ ABCs, which are phenotypically and functionally overlapping with DN2, may exhibit a higher degree of clonal overlap with PB/PCs than with GCBs or with CD11c^-^ subsets. We therefore visualized overlapping clones from DKO female and Yaa-DKO male mice as strings and in Venn Diagrams (Fig. 3G-H; S3E-J). These visualizations revealed a high level of clonal overlap across all subsets, including hundreds of clones that were present in all the subsets. To quantify and compare the level of overlap between the different subsets, we compared the Jaccard index, in which each clone is only counted once in each subset, and the Cosine similarity, in which clone size is also taken into account (Fig. 3I; S3K-J). Neither measure revealed a consistent pattern of similarity with respect to CD11c status. However, ABCs were consistently most highly associated with PB/PCs.

To further verify the relationships between ABCs and the other effector populations in DKOs, we crossed them with Tbet-zsGreen-T2A-CreER^T^^2^-Rosa26-loxP-STOP-loxP-tdTomato mice (termed ZTCE-DKO), where cells expressing T-bet co-express zsGreen and can be traced with Tamoxifen-inducible tdTomato expression^36^. These mice allow for the detection of stable T-bet-expressing cells (zsGreen^+^tdTomato^+^) and cells that previously but no longer express T-bet (zsGreen^-^tdTomato^+^)^36^. ABCs expressed high levels of both ZsGreen and tdTomato, indicating stable expression of T-bet (Fig. 3J-K). CD11c^+^ and CD11c^-^ GC B cells contained both zsGreen^+^tdTomato^+^ and zsGreen^-^tdTomato^+^ although the CD11c^-^ subset was preferentially zsGreen^-^tdTomato^+^ suggesting that both populations can originate from T-bet expressing cells but that CD11c^+^ GC B cells include a greater fraction of stable T-bet expressors (Fig. 3J-K). Although no longer expressing ZsGreen, a substantial fraction of CD11c^+^ PB/PCs and CD11c^-^ PB/PCs were tdTomato^+^, suggesting that both these populations can derive from T-bet-expressing B cells (Fig. 3J-K). In combination with our clonal overlap analyses, these findings suggest that, in this autoimmune setting, ABCs share lineage relationships with both CD11c^+^ and CD11c^-^ GC B cell and PB/PC populations.

### Enrichment for an ISG signature differentiates ABCs from DKO females and males

While ABCs from female and male DKOs exhibited similar BCR repertoire features, their marked differences in function and differentiation suggested that they might employ distinct molecular programs. To gain insights into these mechanisms, we compared their transcriptome by RNA-seq (Fig. 4A). Gene set enrichment (GSEA) and CPDB pathway analyses revealed that ABCs from DKO females were enriched for pathways related to SLE pathogenesis, interferon (IFN) responses, and TLR and complement cascades (Fig. 4B; S4A-C). In line with the known upregulation of both Type I and Type II IFN signatures in SLE patients, female ABCs were enriched for IFN*α* and IFN*γ* responses and upregulated several IFN stimulated genes (ISGs) expressed in SLE PBMCs (Fig. 4B; S4B)^37^. Male ABCs were instead enriched for pathways related to RhoGTPase signaling and platelet activation (Fig. 4C; S4D-E). Given the profound effects of *Tlr7* duplication on the ABCs of DKO males, we next sorted ABCs from Yaa-DKO males and compared their transcriptome to that of ABCs from DKO males (Fig. 4D). Similar to what was observed in female ABCs, the top pathways upregulated in ABCs from Yaa-DKO males were those related to IFN responses (Fig. 4E; S4F-G). ABCs from DKO males were instead enriched for genesets related to hemostasis and platelet activation (S4H-I). Thus, enrichment for an ISG signature differentiates ABCs from DKO females and males and *Tlr7* duplication promotes the upregulation of ISGs in male ABCs.

**Figure 4.**
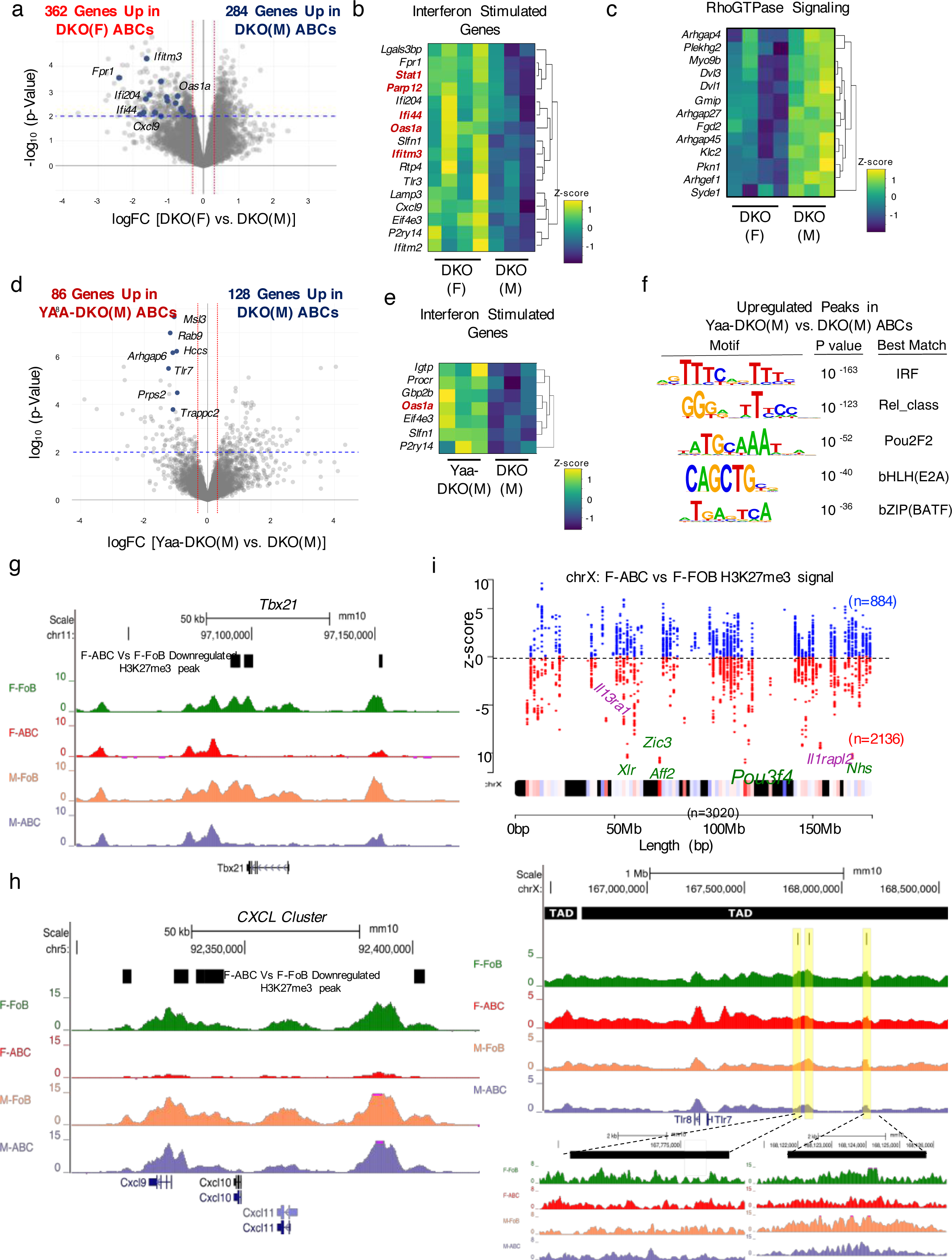
DKO ABCs upregulate ISGs in a sex-specific manner. (A-E) RNA-seq analyses were performed on sorted ABCs (B220^+^CD19^+^CD11c^+^CD11b^+^) from female (F) DKO, male (M) DKO, and male Yaa-DKO mice. (A) Volcano plot showing differentially expressed genes (p <0.01) in ABCs from DKO(F) and DKO(M) mice. Genes in blue show Interferon Stimulated Genes (ISGs) represented in Fig. 4B. (B) Heatmap showing the differential expression of ISGs in ABCs from DKO(F) and DKO(M) mice. (C) Heatmap showing differentially expressed genes in the REACTOME_RhoGTPase_CYCLE geneset. (D) Volcano plot showing differentially expressed genes (p <0.01) in ABCs from Yaa-DKO(M) mice and DKO(M) mice. Genes in blue show genes on the Yaa translocation. (E) Heatmap showing the differential expression of ISGs in ABCs from Yaa-DKO(M) and DKO(M) mice. (F) ATAC-seq was performed on ABCs from DKO(F), DKO(M), and Yaa-DKO(M) mice. Motif enrichment analysis in ATAC-seq peaks that are significantly upregulated (logFC >1.5; p <0.05) in ABCs from Yaa-DKO(M) as compared to DKO(M) mice. (G-I) CUT&RUN for H3K27me3 marks was performed on ABCs from DKO(F) and DKO(M) mice. (G-H) Representative UCSC genome browser tracks of H3K27me3 marks for *Tbx21* gene *(G)* and *CXCL* gene cluster (*H; Cxcl9, Cxcl10 & Cxcl11*). The black boxes in each of the browser track represents genome co-ordinates of significantly down-regulated H3K27me3 peaks F-ABCvsF-FoB (2-fold down and =<0.05 FDR). (I) X-chromosome map of 3092 H3K27me3 peaks across the length the chromosome where each dot represents a H3K27me3 peak. Black bands on X-chromosome are devoid of H3K27me3 signal and the color spectrum represents various H3K27me3 peaks. The z-score (X-chromosome inactivation score) is a DESeq2 Wald statistic showing gain or loss of H3K27me3 signal across 3092 peaks. A total of 2135 peaks had negative z-scores (downregulated) in F-ABC over F-FoB and 884 had positive z-scores (upregulated). Genome browser track of H3K27me3 signals over the *Tlr7* locus, highlighted in yellow are the down-regulated H3K27me3 peaks in F-ABC. The black boxes on the top represent TAD boundaries on the X-chromosome called by machine-learning program Peakachu ^88^ obtained from a murine B cell lymphoma CH12 ^89^.

We next employed ATAC-seq to investigate the chromatin landscape of ABCs derived from the different DKOs. We identified at least 85,000 peaks in ABCs from female, male, and Yaa-DKO male mice (Fig. S4J; Table S1-3). *Tlr7* overexpression induced sufficient changes in the chromatin landscape to enable a motif analysis of the differentially accessible regions (DAR) of Yaa-DKO male ABCs versus male ABCs. Peaks upregulated in ABCs from Yaa-DKO males were enriched for motifs known to be bound by IRFs and NF*κ*B family members (Fig. 4F). Peaks upregulated in ABCs from DKO males were instead enriched for ETS binding sites (Fig. S4K). Consistent with the transcriptional profiles and the enrichment in IRF binding motifs, loci that were differentially accessible in ABCs from Yaa-DKO males as compared to DKO males included a number of ISGs like *Cxcl11*, *Ifi44*, and *Ifitm3* (Fig. S4L). Thus, differences in ISG expression by ABCs are accompanied by a differential enrichment for IRF binding motifs.

We hypothesized that the aberrant gain of an IFN signature by ABCs in females was linked to intrinsic alterations in the epigenetic signatures of these cells and was not simply due to exposure to an IFN-rich environment. To address this hypothesis, we employed the CUT&RUN technique to compare the global loss of a repressive chromatin mark, H3K27me3, at key regulatory loci in ABC and FoB populations sorted from the same mice. Principal component analysis (PCA) of H3K27me3 peaks in ABCs and FOBs from DKO females and males resulted in distinct clustering of samples based on sex and B cell subsets (Fig. S4M). Genome-wide comparison of ABCs and FOBs from DKO females showed distinct loss of H3K27me3 signals at several loci among which *Tbx21,* the classical ABC marker, showed strongest loss in F-ABCs (Fig. 4G; S4N-P). ABCs from DKO males demonstrated a similar loss of H3K27me3 signals at *Tbx21* when compared to FOBs (Fig. 4G). Notably, however, only ABCs from females but not those from males showed selective loss of H3K27me3 marks at ISGs like the CXCL cluster (Fig. 4H; Table S4) suggesting that ABCs in DKO females have intrinsic alterations in their epigenetic signatures that can predispose them to aberrantly gain an IFN signature.

Given that loss of H3K27me3 signals has been shown to accompany XCI, we next specifically investigated the H3K27me3 peaks on the X-chromosome of ABCs and FOBs from DKO females. To gain insights into distinct regulation at various loci on the X-chromosome, we plotted Wald’s z-statistic (X-chromosome deactivation score) obtained from DEseq2 analysis on the 3092 H3K27me3 peaks on the X-chromosomes between ABCs and FoBs from DKO females. H3K27me3 X-chromosome deactivation score plot showed distinct loss of H3K27me3 at over 2092 loci in ABCs as compared to FoBs that also included regulatory loci surrounding the *Tlr7* gene (Fig. 4I). A closer look at the *Tlr7* locus showed that there was a total of 3 peaks (chrX:167770493-167776995, chrX:167827673-167830728 and chrX:168121697-168125788) within the *Tlr7* neighborhood that were deactivated (Fig. 4I). The deactivated loci were located far away from the *Tlr7* gene body but remained in the same topologically associated domains (TADs) suggesting selective deactivation of repressive H3K27me3 marks within the intra-TAD boundaries of the *Tlr7* gene locus. These results suggest that selective loss of repressive chromatin marks in ABCs of DKO females can contribute to the increased expression of X-chromosome linked genes like *Tlr7*.

### Aberrant expansion of CD11c-expressing pre-GC B cells and PBs in DKO females

In addition to ABCs, populations of CD11c-expressing GC B cells and PB/PCs also accumulate to a greater extent in DKO females than DKO males suggesting that they also contribute to the sex-bias in disease development. Since the phenotypic and molecular characteristics of these populations are largely unknown, we investigated them in greater detail. CD11c^+^ GC B cells shared several phenotypic features with ABCs including high expression of T-bet, Fcrl5, and Cxcr3 (Fig. S5A). As compared to CD11c^-^ GC B cells, moreover, the transcriptome of sorted CD11c^+^ GC B cells was enriched for ABC genesets and for IFN responses (Fig. 5A-B; S5B-C). Despite expressing comparable transcript levels of several classical GC target genes, including *Bcl6, Irf8,* and *Spib*, the levels of BCL6 protein were lower in the CD11c^+^ than in the CD11c^-^ populations (Fig. 5C; S5D). In line with the notion that intermediate expression of BCL6 protein in B cells has been associated with a pre-GC B cell state^38^, the CD11c^+^ population contained increased frequencies of BCL6^mid^IRF4^+^ B cells, a profile associated with pre-GC B cells, and was enriched in a pre-GC B cell signature by GSEA (Fig. 5D-E). BCR signaling, as monitored by the phosphorylation of SYK and LYN, was furthermore significantly higher in CD11c^+^ than CD11c^-^ GL7^+^Fas^+^ B cells (Fig. 5F). The CD11c^+^ subset furthermore was less proliferative than CD11c^-^ GL7^+^Fas^+^ B cells (Fig. 5G-H). In contrast, the CD11c^+^ population upregulated pathways related to migration and apoptotic cell clearance and expressed high levels of MerTK, a critical efferocytic receptor (Fig. 5I-J). A greater percentage of CD11c^+^ GL7^+^Fas^+^ B cells than CD11c^-^ GL7^+^Fas^+^ B cells furthermore could engulf apoptotic thymocytes and this was coupled with upregulation of surface MHC-II expression (Fig. S5E-F). Thus, the CD11c^+^ GL7^+^Fas^+^ B cells that aberrantly expand in DKO females likely represent ABCs that have acquired a pre-GC B cell phenotype and can both engulf and present apoptotic debris, thus potentially augmenting autoreactive responses.

**Figure 5.**
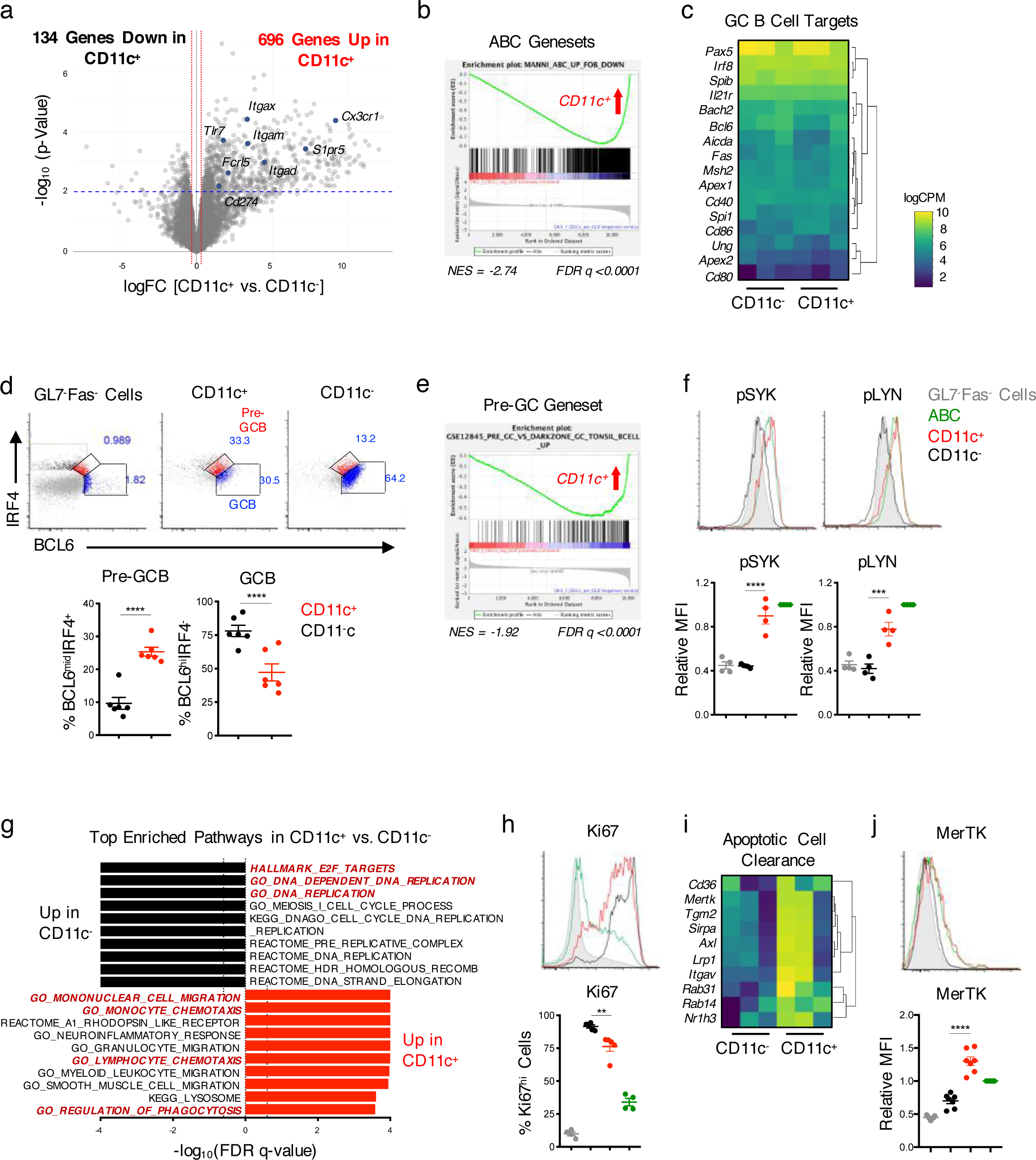
CD11c^+^ GC B cells accumulating in DKO females exhibit a pre-GC B cell phenotype. RNA-seq was performed on sorted CD11c^+^ and CD11c^-^ GL7^+^CD38^lo^ B cells (gated on CD19^+^ splenocytes) from DKO(F) mice. (A) Volcano plot showing genes differentially expressed (p <0.01) between CD11c^+^ CD19^+^GL7^+^CD38^lo^ (CD11c^+^) and CD11c^-^ CD19^+^GL7^+^CD38^lo^ (CD11c^-^) cells. (B) Plot showing the enrichment of the ABC geneset from DKO mice in CD11c^+^ CD19^+^GL7^+^CD38^lo^ cells. (C) Heatmap showing the expression of GC B cell target genes in CD11c^+^ and CD11c^-^ CD19^+^GL7^+^CD38^lo^ cells. (D) Representative plots and quantifications of pre-GC B cells (BCL6^mid^IRF4^+^) and GC B cells (BCL6^hi^IRF4^-^) among CD11c^+^ CD19^+^GL7^+^Fas^+^ (CD11c^+^) and CD11c^-^ CD19^+^GL7^+^Fas^+^ (CD11c^-^) cells. Data representative of and/or pooled from 6 DKO(F) mice and show mean +/- SEM; p-value by paired two-tailed t-tests. (E) Plot showing the enrichment of a pre-GC geneset (GSE12845) in CD11c^+^ CD19^+^GL7^+^CD38^lo^ cells. (F) Representative histograms and quantifications of the phosphorylation of SYK(Y352) and LYN(Y416) in CD11c^+^ CD19^+^GL7^+^Fas^+^ cells (CD11c^+^) (*red*), CD11c^-^ CD19^+^GL7^+^Fas^+^ (CD11c^-^) cells (*black*), and CD19^+^CD11c^+^CD11b^+^ (ABCs) (*green*) from DKO(F) mice. CD19^+^GL7^-^Fas^-^ cells (*gray*) are shown as control. Data representative of and/or pooled from 4 mice and show mean +/- SEM; p-value by 1-way ANOVA followed by Tukey’s test for multiple comparisons. (G) Plot showing the top pathways upregulated in CD11c^-^ CD19^+^GL7^+^CD38^lo^ cells *(black)* and CD11c^+^ CD19^+^GL7^+^CD38^lo^ cells *(red)* by GSEA. Dotted line indicates significance threshold at FDR q <0.25. (H) Representative histogram and quantification of Ki67 expression in the indicated populations from DKO(F) mice. Data representative of and/or pooled from 5 mice and shows mean +/- SEM; p-value by 1-way ANOVA followed by Tukey’s test for multiple comparisons. (I) Heatmap showing the expression of genes related to apoptotic cell clearance in CD11c^+^ and CD11c^-^ CD19^+^GL7^+^CD38^lo^ cells from DKO(F) mice. (J) Representative histogram and quantification of MerTK expression on the indicated populations from DKO(F) mice. Data representative of and/or pooled from 7 mice and show mean +/- SEM; p-value by 1-way ANOVA followed by Tukey’s test for multiple comparisons. *p <0.05, **p <0.01, ***p <0.001, ****p <0.0001.

To gain insights into the molecular profiles of the CD11c^+^ PB/PCs that also aberrantly expand in DKO females, we sorted CD11c^+^ and CD11c^-^ PB/PCs from DKO females and compared their transcriptome by RNA-seq (Fig. 6A). Consistent with their expression of CD11c, CD11c^+^ PB/PCs were enriched for ABC signatures and IFN responses (Fig. 6B; S6A-C). CD11c^+^ PB/PCs furthermore expressed higher surface levels of ABC markers like Cxcr3 and Fcrl5 than CD11c^-^ PB/PCs, although, consistent with the fate mapping studies, they expressed only low levels of T-bet (Fig. 6C). Thus, CD11c^+^ PB/PCs share several transcriptional and phenotypic similarities with ABCs despite downregulating T-bet expression. CD11c^+^ and CD11c^-^ PB/PCs expressed similar transcript levels of *Prmd1* and *Irf4* although the transcriptional programs normally regulated by Blimp1 and Irf4 in PB/PCs were enriched to a greater extent in CD11c^-^ PB/PCs than in CD11c^+^ PB/PCs (Fig. 6D; S6D). CD11c^+^ PB/PCs were more proliferative than CD11c^-^ PB/PCs and expressed higher levels of B220, MHC-II, and *Ciita* (Fig. 6E-F). As compared to CD11c^-^ PB/PCs, CD11c^+^ PB/PCs upregulated pathways related to migration including chemokine receptors like *Ccr3* as well as the expression of pro-inflammatory cytokines such as *Tnf* and *Il1b* and of inflammasome components like *Nlrp3* (Fig. 6H-J). Taken together these data thus suggest that CD11c^+^ PB/PCs represent a population of PBs with distinctive migratory and pro-inflammatory characteristics.

**Figure 6.**
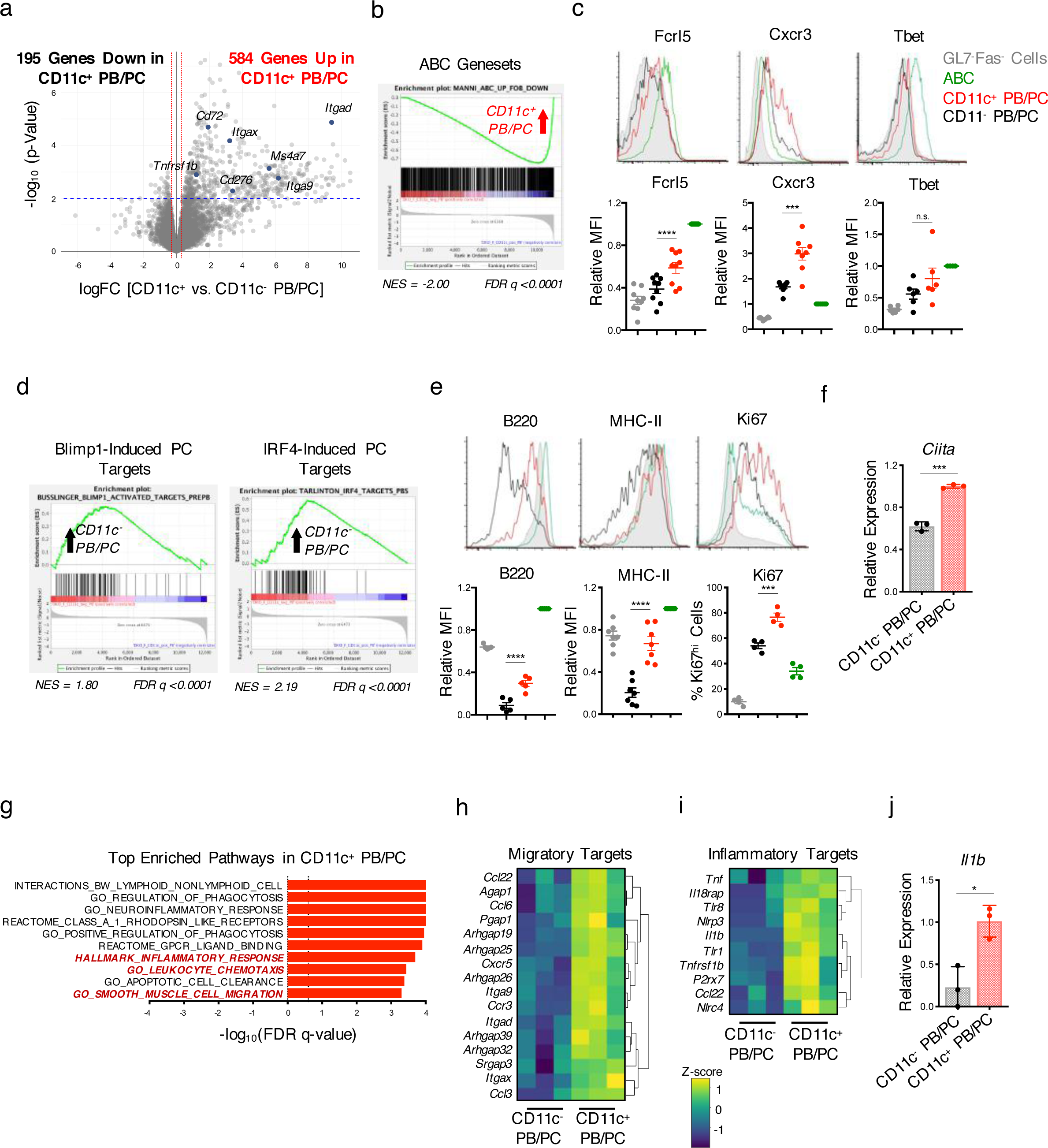
Expansion of CD11c^+^ PBs in DKO females. (A) RNA-seq was performed on sorted CD11c^+^ and CD11c^-^ PB/PCs (CD138^+^TACI^+^) from DKO(F) mice. Volcano plot showing differentially expressed genes (p <0.01) between CD11c^+^ and CD11c^-^ PB/PCs. Blue dots indicate ABC target genes upregulated in CD11c^+^ PB/PCs. (B) Plot showing the enrichment of the ABC geneset from DKO mice in CD11c^+^ PB/PCs. (C) Representative histograms and quantifications of Fcrl5, Cxcr3, and Tbet in CD11c^+^ PB/PCs (CD19^mid/lo^CD138^+^; *red*), CD11c^-^ PB/PCs (*black*), and CD19^+^CD11c^+^CD11b^+^ (ABCs) (*green*) from DKO(F) mice. CD19^+^GL7^-^Fas^-^ cells (*grey*) are shown as a control. Data show mean +/- SEM; *n*>=6; p-value by 1-way ANOVA followed by Tukey’s test for multiple comparisons. (D) Plots showing the enrichment of Blimp1- induced^74^ and IRF4-induced target genes in CD11c^-^ PB/PCs by GSEA. (E) Representative histograms and quantifications of B220, MHC-II, and Ki67 expression in the indicated populations. Data representative of and/or pooled from at least 4 mice and show mean +/- SEM; p-value by 1-way ANOVA followed by Tukey’s test for multiple comparisons. (F) Representative RT-qPCR showing *Ciita* expression in CD11c^+^ or CD11c^-^ PB/PCs sorted from DKO(F) mice as in Fig. 6A. Data representative of 3 independent experiments and show mean +/- SD; p-value by unpaired two-tailed t-test. (G) Plot showing the top pathways enriched in CD11c^+^ PB/PCs as compared to CD11c^-^ PB/PCs by GSEA. Dotted line indicates significance threshold at FDR q<0.25. (H-I) Heatmap showing the expression of migratory *(H)* and inflammatory target genes *(I)* that are differentially expressed (p <0.01) between CD11c^+^ and CD11c^-^ PB/PCs in DKO(F) mice. (J) Representative RT-qPCR showing *Il1b* expression in CD11c^+^ or CD11c^-^ PB/PCs sorted from DKO(F) mice as in Fig. 6A. Data representative of 3 independent experiments and show mean +/- SD; p-value by unpaired two-tailed t-test. *p <0.05, **p <0.01, ***p <0.001, ****p < 0.0001.

### IRF8 and IRF5 cooperate in promoting the generation and differentiation of ABCs in DKO females

The finding that a substantial proportion of effector B cell populations were related to ABCs and previously expressed T-bet prompted us to examine the role of T-bet in the accumulation of ABCs and their progeny in DKO females. We thus generated CD23-Cre *Tbx21*^flox/flox^.DKO mice to specifically delete T-bet in B cells from DKO mice. Despite successful T-bet deletion, lack of B-cell T-bet did not significantly decrease the formation of ABCs as assessed by staining with CD11c and CD11b and other ABC markers such as Fcrl5 and Cxcr3 (Fig. 7A-C). Lack of B cell T-bet also did not affect the accumulation of total or CD11c^+^ GL7^+^Fas^+^ B cells and PB/PCs or the TFH/TFR ratio (Fig. 7D-H). Lack of B cell T-bet did, however, result in a profound decrease in anti-dsDNA IgG2c without a corresponding increase in anti-dsDNA IgG1 (Fig. 7I). Thus, in autoimmune-prone DKO females, B cell T-bet is not necessary for ABC generation or differentiation but is specifically required for the production of IgG2c autoantibodies.

**Figure 7.**
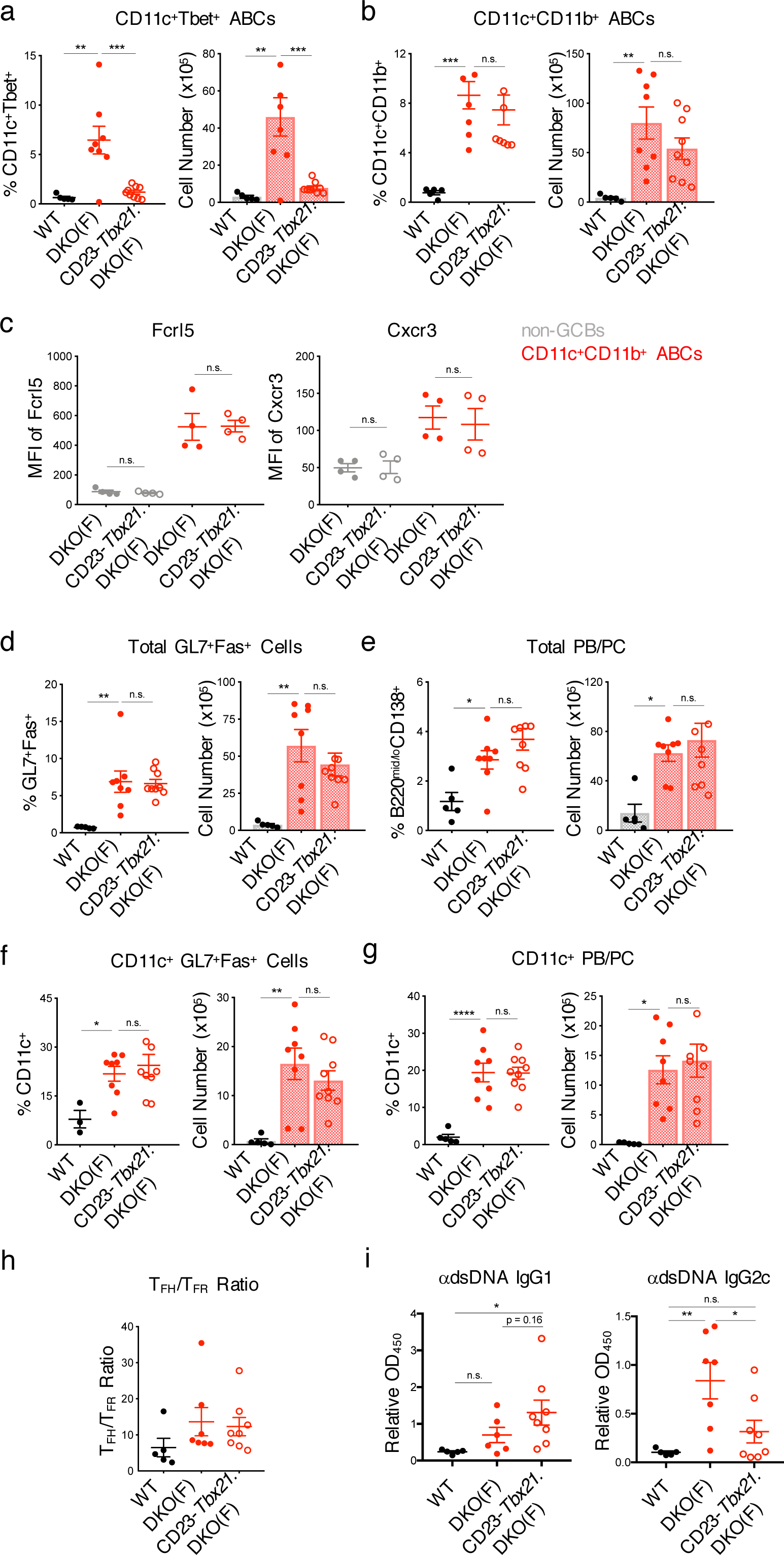
Tbet is not required for ABC accumulation in DKO mice. (A-B) Quantifications showing the numbers of CD11c^+^Tbet^+^ ABCs *(A)* and CD11c^+^CD11b^+^ ABCs *(B)* from WT, DKO(F), and CD23-Cre.*Tbx21*^f/f^.DKO (CD23-*Tbx21*.DKO(F)) mice. Data pooled from at least 6 mice per genotype and show mean +/- SEM; p-value by 1-way ANOVA followed by Tukey’s test for multiple comparisons. (C) Plots showing the MFI of Fcrl5 and Cxcr3 on the surface of CD11c^+^CD11b^+^ ABCs *(red)* from the indicated mice. Non-GCBs (CD19^+^GL7^-^Fas^-^; *gray*) are shown as control. Data pooled from 4 mice per genotype and show mean +/- SEM; p-value by unpaired two-tailed t-tests. (D-G) Quantifications showing the numbers of total GL7^+^Fas^+^ B cells *(D)*, total PB/PCs (*E)*, CD11c^+^ GL7^+^Fas^+^ B cells *(F)*, and CD11c^+^ PB/PCs *(G)* from the indicated mice. Data pooled from at least 6 mice per genotype and show mean +/- SEM; p-value by 1-way ANOVA followed by Tukey’s test for multiple comparisons. (H) Quantifications showing the ratio of T_FH_ to T_FR_ cells ratio from the indicated mice. Data pooled from 5 mice per genotype and show mean +/- SEM; p-value by 1-way ANOVA followed by Tukey’s test for multiple comparisons. (I) Pooled ELISA data of anti-dsDNA IgG1 and IgG2c levels in the serum from the indicated mice. Data pooled from at least 6 mice per genotype and show mean +/- SEM; p-value by 1-way ANOVA followed by Tukey’s test for multiple comparisons. * p <0.05, ** p <0.01, *** p <0.001, **** p <0.0001.

Given that global IRF5 deletion profoundly decreased ABC formation in DKO females ^20, 39^, we next employed a similar strategy to investigate whether B-cell expression of IRF5 was specifically required for ABC generation and differentiation. Since IRF8 was also identified as a potential upstream regulator of ABCs in DKO females versus males (Fig. S7A), we also extended this analysis to B cell IRF8. ABC accumulation was significantly decreased in CD23- Cre.*Irf5^flox/+^*DKO, CD23-Cre.*Irf5^flox/flox^*DKO, and CD23-Cre.*Irf8^flox/flox^*DKO females (Fig. 8A-B). Lack of B-cell IRF5 and IRF8 also markedly affected the expansion of GC B cells and CD11c^+^ PB/PCs but only the absence of IRF5 affected the accumulation of total PB/PCs (Fig. 8C-F). Absence of B cell IRF5 or IRF8 also significantly impacted T_FH_ responses and the production of anti-dsDNA IgG2c antibodies (Fig. 8G; S7B-C). To delineate the relative contributions of IRF5 and IRF8 to the generation of ABCs, we utilized an *in vitro* culture system^20^. Lack of either IRF5 or IRF8 reduced the ability of DKO B cells to differentiate into ABCs albeit not in an identical manner, since deletion of IRF8 impaired CD11c upregulation while lack of IRF5 diminished T-bet induction (Fig. 8H-I). Both IRF5 and IRF8 were required for the enhanced production of CXCL10 by DKO ABCs and ChIP-qPCR demonstrated increased binding of both IRF5, as we previously reported^20^, and IRF8 to the regulatory regions controlling the *Cxcl* cluster of genes in DKO B cells (Fig. 8J-K). Deleting IRF8 in DKO B cells impaired the ability of IRF5 to bind to these regions and lack of IRF5 decreased binding of IRF8 to these sites (Fig. 8K). Taken together these data support the notion that cooperation between IRF5 and IRF8 promotes the aberrant accumulation, function, and differentiation of ABCs in DKO females.

**Figure 8.**
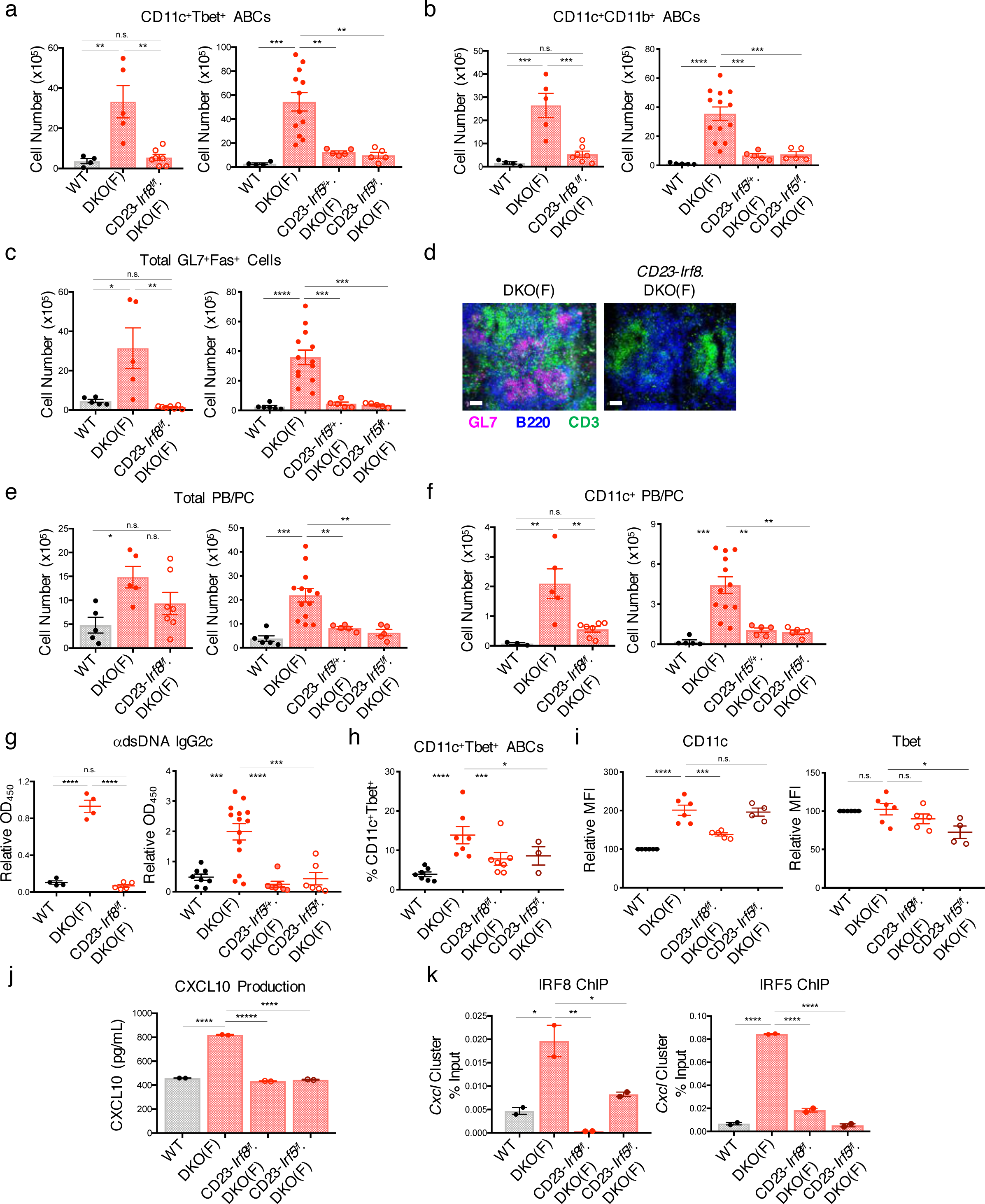
IRF8 and IRF5 are required for ABC generation and differentiation in DKO females. (A-C) Quantifications showing the numbers of CD11c^+^Tbet^+^ ABCs *(A)*, CD11c^+^CD11b^+^ ABCs *(B)*, and total GL7^+^Fas^+^ B cells *(C)* from the spleens of WT, DKO(F), CD23-*Irf8*.DKO(F), CD23-*Irf5^f/+^*.DKO(F), and CD23-*Irf5.*DKO(F) mice. Data pooled from at least 5 mice per genotype and show mean +/- SEM; p-value by 1-way ANOVA followed by Tukey’s test for multiple comparisons. (D) Representative immunofluorescence images of B220 *(blue),* GL7 *(pink)*, and CD3 *(green)* on the indicated mice. Data representative of at least 4 mice per genotype and show mean +/- SEM; p-value by unpaired two-tailed t-test. (E-F) Quantifications showing the numbers of total PB/PCs (B220^mid/lo^CD138^+^) *(E)* and CD11c^+^ PB/PCs *(F)* from the indicated mice. Data pooled from at least 5 mice per genotype and show mean +/- SEM; p-value by 1-way ANOVA followed by Tukey’s test for multiple comparisons. (G) ELISA data showing anti-dsDNA IgG2c in the serum from the indicated mice. Data pooled from at least 4 mice per genotype and show mean +/- SEM; p-value by 1-way ANOVA followed by Tukey’s test for multiple comparisons. (H-K) CD23^+^ B cells from the indicated mice were stimulated for 2-3d with *α*IgM, *α*CD40, and IL-21. (H) Quantification of CD11c^+^Tbet^+^ B cells after 3d culture. Data pooled from at least 3 mice per genotype and show mean +/- SEM; p-value by 1-way ANOVA followed by Tukey’s test for multiple comparisons. (I) Quantifications showing the expression of CD11c and Tbet in the indicated strains after 3d culture. Data pooled from at least 4 mice per genotype and show mean +/- SEM; p-value by 1-way ANOVA followed by Tukey’s test for multiple comparisons. (J) ELISA data showing CXCL10 production in the supernatants after 3d culture. Data representative of 3 independent experiments and show mean +/- SD; p-value by 1-way ANOVA followed by Tukey’s test for multiple comparisons. (K) Representative ChIP-qPCR of IRF8 and IRF5 binding at the *Cxcl* cluster in the indicated mice. Data representative of 3 independent experiments and show mean +/- SD; p-value by 1-way ANOVA followed by Tukey’s test for multiple comparisons. *p <0.05, **p <0.01, ***p <0.001, ****p <0.0001.

### TLR7 duplication in DKO males promotes ABC dissemination and severe immunopathogenesis

While SLE preferentially affects females, males with lupus often exhibit a more rapid and severe course. Similarly, the shorter survival of Yaa-DKO males than DKO females suggested a more severe immunopathogenesis. Interestingly, the early mortality and renal damage of Yaa-DKO males was accompanied by greater frequencies of ABCs in the blood and kidneys (Fig. 9A-C; S8A-B). Furthermore, a histopathological analysis revealed that Yaa-DKO males also exhibited prominent inflammatory infiltrates in the lungs although other organs, like the colon and the pancreas, were unaffected (Fig. 9D; S8C). FACS analysis demonstrated that the lungs of Yaa-DKO males displayed a marked accumulation of ABCs, CD11c^+^ and CD11c^-^ GC B cells, and CD11c^+^ and CD11c^-^ PB/PCs as well as activated T cells (Fig. 9E; S8D). DKO females also exhibited lung inflammation, but as compared to age-matched Yaa-DKO males, the findings were less severe (Fig. 9D). Lung infiltrates in DKO females, however, worsened with age and were markedly ameliorated in aged CD23-Cre.*Irf8^flox/flox^*DKO females suggesting a crucial contribution of ABCs to the pulmonary inflammation (Fig. 9F). Thus, *Tlr7*-driven expansion of ABCs and their progeny can promote their accumulation in the lungs and the development of severe pulmonary inflammation.

**Figure 9.**
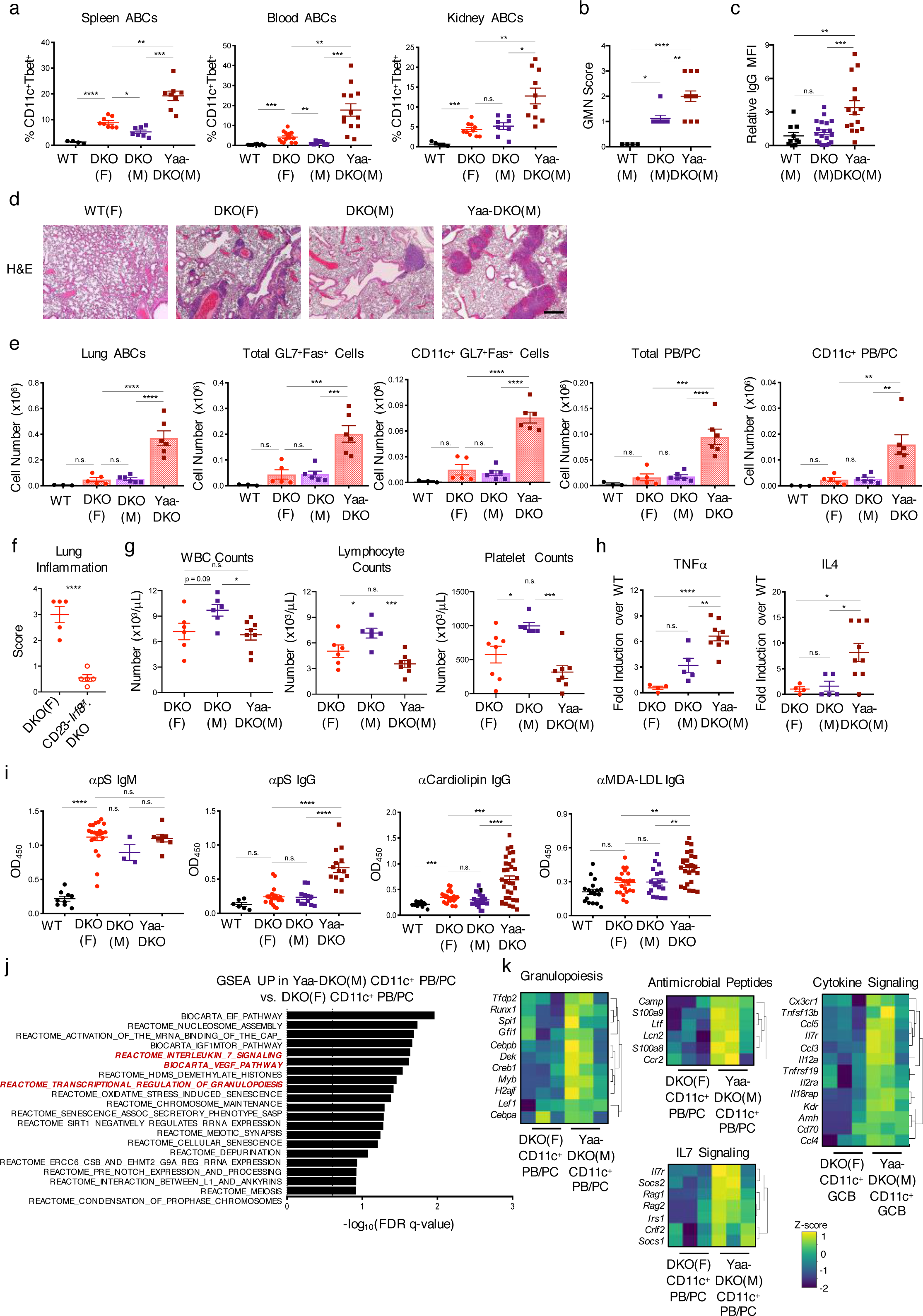
Duplication of *Tlr7* in DKO males promotes ABC dissemination and severe immunopathogenesis. (A) Quantification of CD11c^+^Tbet^+^ ABCs from the spleens, blood, and kidneys of WT, DKO(F), DKO(M), and Yaa-DKO(M) mice. Data shows mean +/- SEM; *n*>=5 mice per genotype; p-value by 1-way ANOVA followed by Tukey’s test for multiple comparisons. (B) Glomerulonephritis (GMN) scores of kidneys from the indicated mice. Data shows mean +/- SEM; *n>*=4 per genotype; p-value by 1-way ANOVA followed by Tukey’s test for multiple comparisons. (C) Quantification of IgG deposition in the kidneys from the indicated mice. Data shows mean +/- SEM; *n*>=10 fields across 2-3 mice per genotype; p-value by 1-way ANOVA followed by Tukey’s test for multiple comparisons. (D) Representative H&E images of lungs from the indicated mice. Data representative of 2 WT, 4 DKO(F), 4 DKO(M), and 4 Yaa-DKO(M) mice (30-36wk). (E) Quantifications of the numbers of CD11c^+^CD11b^+^ ABCs, total GL7^+^Fas^+^ B cells, CD11c^+^ GL7^+^Fas^+^ B cells, total PB/PCs, and CD11c^+^ PB/PCs in the lungs from the indicated mice. Data show mean +/- SEM; *n*>= 3 mice per genotype; p-value by 1-way ANOVA followed by Tukey’s test for multiple comparisons. (F) Plot showing the inflammation score in the lungs of 45wk-old CD23-Cre.*Irf8*^f/f^.DKO(F) and *Irf8*^f/f^.DKO(F) control mice as determined by H&E staining. Data shows mean +/- SEM; *n*=5 per genotype; p-value by unpaired two-tailed t-test. (G) Plots showing the numbers of white blood cells, lymphocytes, and platelets in the blood from the indicated mice. Data show mean +/- SEM; *n*>=6 per genotype; p-value by 1-way ANOVA followed by Tukey’s test for multiple comparisons. (H) Luminex data showing the fold-increase in serum levels of TNF*α* and IL-4 in DKO(F), DKO(M), and Yaa-DKO(M) mice relative to WT. Data pooled from at least 4 mice per genotype and show mean +/- SEM; p-value by 1-way ANOVA followed by Tukey’s test for multiple comparisons. (I) ELISA data for anti-phosphatidylserine (pS) IgM and IgG, anti-Cardiolipin IgG, and anti-MDA-LDL IgG in the serum from the indicated mice. Data shows mean +/- SEM; *n*>=3 per genotype; p-value by 1-way ANOVA followed by Tukey’s test for multiple comparisons. (J) RNA-seq was performed on sorted CD11c^+^ PB/PCs (CD138^+^TACI^+^) from Yaa-DKO(M) mice as in Fig. 6A. Plot shows top pathways enriched in CD11c^+^ PB/PCs from Yaa-DKO(M) mice as compared to CD11c^+^ PB/PCs from DKO(F) mice as determined by GSEA. Dotted line indicates significance threshold at FDR q <0.25. (K) Heatmaps showing the expression of genes enriching the granulopoiesis, antimicrobial peptide, cytokine signaling, and IL7 signaling genesets in Yaa-DKO CD11c^+^ PB/PCs. *p <0.05, **p <0.01, ***p <0.001, ****p <0.0001.

Given that the marked lung inflammation in the setting of TLR7 dysregulation was reminiscent of the pathophysiology not only of SLE but also of severe viral infections such as COVID-19, we conducted additional hematologic and serologic analyses to assess whether other parameters known to be altered in this infection were similarly affected. A peripheral blood count demonstrated lower lymphocyte and platelet counts but increased monocytes in Yaa-DKO males than DKO males (Fig 9G; S8E). Yaa-DKO males also exhibited elevated levels of serum TNF*α* and IL4 (Fig. 9H). We also assessed the production of antiphospholipid antibodies. While all DKOs produced higher levels of anti-phosphatidylserine (pS) IgM antibodies than WT controls, only Yaa-DKO males produced anti-pS IgG, anti-cardiolipin, and anti-MDA-LDL IgG (Fig. 9I). In line with the known ability of ABCs to produce antiphospholipid antibodies in response to pathogens and mediate hematologic abnormalities^40^, anti-pS IgG and anti-cardiolipin antibodies correlated with ABC and PB/PC frequencies and there was a significant inverse association between ABC frequencies and platelet counts (Fig. S8F-G). Thus, *Tlr7* duplication in DKO males results in hematologic and serologic abnormalities that can be associated not only with SLE but also with severe viral infections like COVID-19.

To gain insights into the molecular features that might result in a more rapid and severe TLR7-induced immunopathogenesis in Yaa-DKO males than in DKO females, we compared the transcriptomes of ABCs and their progeny sorted from Yaa-DKO males with those of the corresponding populations sorted from DKO females. Only minimal differences were observed by GSEA between the ABCs of DKO females and those of Yaa-DKO males (Fig. S8H). However, a comparison of CD11c^+^ preGC B cells and CD11c^+^ PBs demonstrated that, as compared to DKO females, the populations derived from Yaa-DKO males were enriched for pathways involved in the production of antimicrobial peptides, cytokine interactions and signaling, and transcriptional regulation of granulopoiesis (Fig. 9J-K; S8I-J). Thus, CD11c^+^ effectors in Yaa-DKO males upregulate pathways that can enhance their pathogenicity and potentially enable their trans-differentiation, as has been recently proposed for PBs in COVID-19 patients^41^.

## DISCUSSION

The mechanisms that underlie the sexual dimorphism observed in responses to infections and vaccinations, and in autoimmune diseases remain incompletely understood. Here, we delineate sex-specific differences in the function and differentiation of ABCs, a subset of B cells that are emerging as critical mediators of antiviral responses and pathogenic players in autoimmunity. Using a spontaneous model of lupus where disease preferentially develops in females, we demonstrate that female ABCs exhibit a greater ability than male ABCs to accumulate, acquire an ISG signature, and further differentiate into effector populations, which include CD11c^+^ pre-GC B cells and CD11c^+^ PBs. BCR sequencing and fate mapping reveal oligoclonal expansion and relatedness amongst ABCs, GC B cells, and PB/PC populations irrespective of the expression of CD11c. Genetic studies demonstrate a critical role for TLR7, IRF5, and IRF8 in promoting these abnormalities in females. Duplication of *Tlr7* in males overrides the sex-bias and triggers severe immunopathology marked by intense lung inflammation and early mortality. Thus, sex-specific differences permeate several aspects of ABC biology in autoimmune settings suggesting that this compartment may be uniquely endowed to function in a sex-specific manner.

Our studies demonstrate marked differences in the ability of female and male DKO ABCs to express an ISG signature, which has recently been shown to be upregulated in ABCs and PCs from SLE patients and is one of the best-known features not only of this disease but also of antiviral responses^7, 42,43,44^. The ISG signature observed in SLE patients indeed overlaps with that detected upon viral infections and immunizations and an inability to upregulate ISGs has recently been shown to distinguish severe from mild-to-moderate COVID-19 patients^45,46,47^. Our epigenetic and genetic analyses furthermore indicate that this system is critically reliant on the dysregulation of IRF5 and IRF8 activity, well-known controllers of ISGs whose variants have long been associated with lupus pathogenesis and whose activity can be a target of viral evasion strategies ^47,48,49^. While we cannot rule out that TLR7-driven IFN*α* production by pDCs could contribute to the differential expression of the ISG signature in DKO females and males, loss of repressive epigenetic markers at ISGs were selectively observed in ABCs, but not FoBs, from DKO females supporting the notion that acquisition of this signature is controlled in large part by cell-intrinsic mechanisms rather than exposure to an IFN-rich environment.

Sex-based differences were also observed in the ability of ABCs to further differentiate into effector subsets. Despite similar T_FH_ responses to DKO males, DKO females exhibited a more robust expansion of GC B cells and PB/PCs, which in addition to classical CD11c^-^ subsets, also included CD11c^+^ expressing subsets. BCR sequencing and fate mapping uncovered surprising relationships of ABCs not only with CD11c^+^ pre-GC B cells and CD11c^+^ PBs, but also with CD11c^-^ GC B and CD11c^-^ PC, which exhibited remarkably different transcriptional profiles from the CD11c^+^ B cell populations. The IRFs may again be a crucial component of this dysregulation. Indeed, the known ability of IRFs to homo/heterodimerize and target ISREs as well as interact with other transactivators like the Ets protein PU.1^48^ may be well-suited to help confer a high degree of heterogeneity to ABCs and their progeny and fine-tune their transcriptional profiles. The pairing of IRF5 with IRF8 in this compartment may provide ABCs and their CD11c^+^ progeny with a more “innate” quality than traditional CD11c^-^ B cell subsets and facilitate their distinctive combination of innate and adaptive functions. Changes in IRF8 activity, like those mediated by

ROCK2 phosphorylation^50^, could instead result in the downregulation of typical ABC transcriptional targets like CD11c and help promote its interaction with other transactivators enabling ABCs to give rise to both CD11c^+^ and CD11c^-^ effector progeny. In support of this notion preliminary studies indicate that TLR7 engagement can inhibit ROCK2 activation thus directly affecting this balance.

In contrast to the marked effects observed upon deletion of IRF5 and IRF8, lack of B-cell T-bet in DKOs exerted more selective effects suggesting that, in this autoimmune setting, the generation and differentiation of CD11c^+^ B cells is less reliant on T-bet. T-bet however played a critical role in the acquisition of functional capabilities, such as the production of anti-dsDNA IgG2a/c antibodies, that have been associated with these cells. Given that a small fraction of FoBs was found to express T-bet in the fate mapping studies, additional analyses will furthermore be required to establish whether these T-bet^+^ cells are destined to acquire a full-fledged ABC phenotype or whether they represent a separate pool of T-bet^+^ B cells that can also differentiate into a heterogenous progeny and are critical for autoantibody production.

VH sequencing demonstrated profound oligoclonal expansion and further confirmed common clonal relationships between ABCs, GC B cells, and PB/PCs, which again could be observed within both CD11c^+^ and CD11c^-^ compartments. Interestingly, different degrees of clonal overlap could be observed between ABCs and different progenies within distinct mice suggesting that while both extrafollicular and GC-like differentiation pathways may be available to these cells, the precise routes employed by ABCs to undergo terminal differentiation can vary depending on the specific inflammatory milieus that they are exposed to. Although the spontaneous GCs observed in this autoimmune setting displayed some atypical features, as evidenced by the finding that these GCs were exquisitely sensitive to the absence of IRF8 unlike those observed upon T-dependent immunization^51^, the ABC-derived GC populations did exhibit higher levels of SHM than ABCs suggesting that these GCs were functional. Notably, several of the VH regions overexpressed in DKO ABCs and their progeny, including VH1 (J558) and VH14 (SM7), have previously been associated with the production of lupus autoAbs and with the expanded ABCs of SLC^-/-^ mice, another spontaneous autoimmune model^52^.

The plasticity of ABCs and a limited capability to acquire GC-like as well as PB phenotypes have been previously observed in *Ehrlichia* and influenza infection models^9, 10, 53^ suggesting that, upon encountering a pathogen, these differentiation pathways are available to ABCs, albeit in a restricted manner. One of the consequences of TLR7 overexpression in our autoimmune model furthermore was the dissemination of ABCs in the blood and their accumulation in the lungs, a pattern that was also transiently exhibited by T-bet^+^ B cells shortly after influenza infection^9^. Strikingly, the lung infiltrates in aged DKO females were markedly ameliorated in mice lacking IRF8 in B cells supporting a key role for ABCs and their progeny in promoting the pulmonary inflammation observed in DKOs. Thus, some of the pathogenesis of SLE may reflect a breakdown in the regulatory mechanisms aimed at restricting the differentiative routes and dissemination of ABCs in time and space. In this regard, the ability of the SWEF proteins to coordinate cytoskeletal organization and IRF function^21^ may be particularly important. Indeed, by ensuring the proper positioning, cell-cell interactions, and transient migration of ABCs and by restricting the accessibility to IRF controlled transcriptional programs, these molecules could help ensure that a rapid initial response is coupled with the transient generation of a pool of progeny, which is diverse but limited in size.

Although lower levels of TLR7 responsiveness by male ABCs may protect DKO males from the development of lupus, the transcriptional profile of male ABCs showed enrichment for Rho GTPase signaling pathways. Dysregulation of these pathways has been linked to well-known disorders such as hypertension and cardiovascular disease, which also show a sexually dimorphic pattern but preferentially affect males^54, 55^. This raises the intriguing possibility that lower levels of TLR7 engagement by male ABCs could not only result in suboptimal antiviral defenses but also promote a distinct set of pathogenic responses marked by vascular inflammation and thrombosis. Surprisingly, TLR7 duplication in Yaa-DKO males resulted in more profound immunopathogenesis than that observed in DKO females suggesting that females may have evolved mechanisms to contain the overwhelming inflammatory effects that may accompany the greater levels of TLR7 stimulation that they are predisposed to due to incomplete XCI. In this regard, it is interesting to note that both our studies and recent work in humans suggest that ABCs may be uniquely susceptible to partially escape XCI ^56^. We cannot, however, rule out that the differences between Yaa-DKO males and DKO females primarily result from complete rather than partial escape from XCI secondary to the translocation of the X-chromosome segment onto the Y-chromosome. While our studies have highlighted a crucial role for TLR7 in mediating the sex-differences in the development of lupus in this model, sex-hormones like estrogen and androgen have also been shown to regulate B cell responses including their positioning and survival and to modulate disease pathogenesis ^57, 58^ and thus are also important contributors to these differences.

Intriguingly, several of our findings are reminiscent of the pathophysiology of COVID-19 where disease outcomes have shown striking age- and sex-dependent differences^1^. Indeed, expansion of T-bet^+^ B cells (including activated naïve and DN) and PBs, which in some cases can exhibit reprogramming potential and lower levels of SWAP-70, have all been observed in severe COVID-19 patients^12, 13, 41, 59^. Some of the features known to accompany disease severity in COVID-19 such as lymphopenia, thrombocytopenia, production of antiphospholipid antibodies, and a broad array of cytokine responses^60, 61^ could be observed in Yaa-DKO males in the absence of any viral infections suggesting that these effects could directly result from unbridled TLR7 stimulation. Whether the reported ability of ABC-like cells to accumulate with age in adipose tissue in an IL-1-dependent manner^62^ could provide a ready depot of these cells and lower the threshold of TLR7 stimulation needed by these cells to acquire full pathogenic functions will be an important question to be addressed given the link between obesity and COVID-19 outcomes. The marked sex-differences in the pathways utilized by female and male ABC subsets suggest that the effectiveness of approaches targeting these cells may demonstrate sex-dependent differences. Such a scenario may be encountered, for instance, in the case of statins or geranylgeranylation inhibitors that can inhibit Rho-GTPases and have vaccine adjuvant properties^63^. Sex-specific differences in ABC function and differentiation could thus not only contribute to the well-known sex-bias that underlies several autoimmune diseases but also broadly impact responses to pathogens and vaccination.

## METHODS

### Mice

DEF6-deficient (*Def6*^tr/tr^) mice were generated by Lexicon Pharmaceuticals, Inc. using a gene trapping strategy as previously described^22^. Swap70-deficient (*Swap70*^-/-^) were generated as previously described^22^. *Def6*^tr/tr^*Swap70*^-/-^ (DKO) mice were generated by crossing *Def6*^tr/tr^ mice with *Swap70*^-/-^ mice that had been backcrossed onto C57BL/6 background for >10 generations^22^. C57BL/6 mice, *Irf8*^f/f^ mice (#014175), and *Tbx21^f/f^* mice (#022741) were from Jackson Laboratory. B6.SB-*Yaa*/J were originally provided by Darryl Roopenian and are available thru Jackson (Yaa; #000483) mice, *Tlr7*-deficient were originally provided by Eric Pamer and are available thru Jackson (*Tlr7^-/-^*; #008380)^64^. Male Yaa mice were crossed with female DKO mice to generate Yaa-DKO male mice^65^. *Irf8^f^*^/f^ mice, *Tbx21^f/f^ mice,* and *Tlr7*^-/-^ mice were crossed with DKO mice to generate *Irf8^f/f^*.DKO, *Tbx21^f/f^*.DKO mice, and *Tlr7*^-/-^.DKO mice. DKO Blimp1-YFP and *Irf5^f/f^* DKO mice were previously described^20, 66^. CD23-cre mice were provided by Jayanta Chaudhuri and were previously described^67^. Tbet-zsGreen-T2A-CreER^T^^2^-Rosa26-loxP-STOP-loxP-tdTomato DKO (ZTCE-DKO) mice were generated by crossing DKO mice with Tbet-zsGreen-T2A-CreER^T^^2^ provided from Jinfang Zhu^36^ and with B6.Cg-*Gt(ROSA)26Sor^tm^*^14^*^(CAG-tdTomato)Hze^*/J (#007914, Jackson). ZTCE-DKO mice were treated with Tamoxifen by oral gavage 3d before experiments were conducted. All mice used in the experiments were kept under specific pathogen-free conditions. All the experiments were carried out following institutional guidelines with protocols approved by the Institutional Animal Care and Use Committee of the Hospital for Special Surgery and WCMC/MSKCC.

### Antibodies and Flow Cytometry

The following monoclonal antibodies to mouse proteins were used for multi-parameter flow cytometry: B220 (RA3-6B2; 400x), CD4 (RM4-5; 400x), CD45 (HI30; 400x), CD11b (M1/70; 400x), CD11c (N418; 400x), CD19 (HIB19; 400x), CD21 (7E9; 200x), CD23 (B3B4; 200x), CD44 (IM7; 200x), CXCR3 (CXCR3-173; 200x), MHC-II (AF6-120.1; 600x), and T-bet (4B10; 800x) were obtained from BioLegend. Streptavidin-conjugated antibodies were also obtained from BioLegend. Antibodies to BCL6 (K112-91; 100x), CD138 (281-2; 1200x), CXCR5 (2G8; 200x), Fas (Jo2; 200x), FcRL5 (509F6; 200x), and GL7 (600x) were obtained from BD. Antibodies to Foxp3 (FJK-16s; 100x), IgD (11-26; 500x), IgM (II/41; 1000x), IRF4 (3E4; 200x), IRF8 (V3GYWCH; 200x), Ki67 (solA15; 800x), MerTK (DS5MMER; 200x), and PD1 (J43; 200x) were obtained from eBioscience. For intracellular staining, cells were fixed after surface staining at 4°C with the Transcription Factor Staining Kit (eBioscience) following the manufacturer’s instructions. For intracellular cytokine staining, splenocytes were stimulated with 50μg/mL PMA and 1μM Ionomycin for 4hr. Cells were incubated with BrefeldinA for the final 3hr of stimulation. After stimulation, cells were fixed and permeabilized with a Transcription Factor Staining Kit (eBioscience) and stained using anti-IFN*γ* (XMG1.2; 200x; BioLegend) and recombinant mouse IL21R Fc Chimera (600x; R&D) followed by PE-labeled affinity-purified F(ab’)2 fragment of goat anti-human Fc*γ* (Jackson ImmunoResearch). For detection of phosphorylated antigens, splenocytes were fixed in BD Fixation Buffer for 20min at RT. Cells were then washed and permeabilized in 90% methanol for 30min at -20°C and then incubated with antibodies against phosphorylated SYK (Y352; Cell Signaling) or LYN (Y416; Cell Signaling) for 45min at room temperature. LysoTracker staining was done before surface staining with 20nM LysoTracker-Deep Red (Thermo) for 45min at 37°C in RPMI 1640. All data were acquired on a BD FACS Canto and analyzed with FlowJo (TreeStar) software.

### Cell Sorting

Single-cell suspensions were prepared from spleens of the indicated mice. For sorting ABCs, splenocytes were pre-enriched for B cells with B220 microbeads (Miltenyi Biotec) following manufacturer’s instructions. B cells were stained with CD11c (N418), CD11b (M1/70), CD19 (HIB19), B220 (RA3-6B2), and CD23 (B3B4) and were sorted on FACS Aria or Influx (BD). ABCs were collected for RNA or cultured for 7d with 1μM Imiquimod in RPMI 1640 medium (Corning) supplemented with 10X FBS (Atlanta Biologicals), 100U/mL Penicillin (Corning), 100mg/mL Streptomycin (Corning), 1X non-essential amino acids (Corning), 2mM L-Glutamine (Corning), 25mM HEPES (pH7.2-7.6; Corning), and 50μM *β*-Mercaptoethanol. For sorting CD11c^+^ and CD11c^-^ GC B-like cells and PB/PCs, splenocytes were pre-enriched using biotinylated antibodies against B220 and CD138 and with streptavidin-conjugated microbeads (Miltenyi Biotec) following manufacturer’s instructions. Cells were stained with CD11c (N418), CD19 (HIB19), CD138 (281-2), Fas (Jo2), GL7, and TACI (8F10) and were sorted on FACS Aria or Influx (BD).

### B Cell Cultures

CD23^+^ B cells were purified from single cell suspensions of splenocytes with biotinylated anti-CD23 (BD Bioscience) and streptavidin microbeads (Miltenyi) according to manufacturer’s instructions. Cells were cultured for 2-3d in RPMI 1640 medium (Corning) supplemented with non-essential amino acids (Corning), 2mM L-Glutamine (Corning), 25mM HEPES (pH 7.2-7.6), and 50μM *β*-Mercaptoethanol and stimulated with 5μg/mL F(ab’)2 anti-mouse IgM (Jackson ImmunoResearch), 5μg/mL purified anti-mouse CD40 (BioXcell), and 50ng/mL IL21 (Peprotech). Thymocyte engulfment assays were performed as previously described^68^. In brief, thymocytes were harvested and isolated from 4-6 week-old WT mice and treated with 50μM Dexamethasone for 4hr to induce apoptosis. Apoptotic thymocytes were then stained with 1μM CypHer5E (GE Healthcare) for 45min at 37°C in serum-free Hank’s Balanced Sodium Solution (HBSS). Stained thymocytes were co-cultured with splenocytes at a 1:10 splenocyte to thymocyte ratio. Apoptotic thymocytes were removed by washing with cold PBS and splenocytes were assessed for efferocytosis by flow cytometry. MHC-II expression was compared on efferocytic (CypHer5E^+^) and non-efferocytic (CypHer5E^-^) B cells.

### Real-time RT-PCR and Chromatin Immunoprecipitation (ChIP) Assays

Total RNA was isolated using the RNeasy Plus Mini kit (Qiagen). cDNAs were prepared using the iScript cDNA synthesis kit (Biorad). Real-time PCR was performed using the iTaq Universal SYBR Green Supermix (Biorad). Gene expression was calculated using the *ΔΔ*Ct method and normalized to *Ppia*. Primers for *Ciita* and *Tlr7* were from Qiagen. Custom primers were used for *Ppia* (F: 5’-TTG CCA TTC CTG GAC CCA AA-3’; R: 5’-ATG GCA CTG GCG GCA GGT CC-3’) and *Il1b*. For ChIP assays, B cell cultures were harvested at d2 and chromatin extracts were prepared using the truChIP Chromatin Shearing Reagent Kit (Covaris). 100μg of sonicated DNA protein complexes were used for immunoprecipitations with anti-IRF5 (Abcam #ab21689), anti-IRF8 (Cell Signaling; #5628), or normal anti-rabbit Ig control antibodies. DNA purified from the immunoprecipitates and inputs was analyzed by RT-qPCR using the following primers: *Cxcl cluster* F 5’-AGT AGT CCC CAC TGT CTG ACT-3’; *Cxcl cluster* R 5’-GTG AGT CCC TTT AGC ACC AGA-3’.

### ELISAs and Antigen Microarrays

For anti-dsDNA ELISA, plates were coated with 100μg/mL salmon sperm DNA (Invitrogen; #AM9680) at 37°C overnight and blocked in 2% BSA in PBS at room temperature for 2hr. For anti-cardiolipin and anti-phosphatidylserine ELISA, Immulon 2HB plates (Thermo) were coated with 75μg/mL of cardiolipin or with 30μg/mL phosphatidylserine dissolved in 100% ethanol overnight. Sera were diluted 1:200 and incubated on coated plates at 25°C for 2hr. Supernatants from sorted ABCs were used neat after 7d culture with 1μM Imiquimod. Plates were then incubated with HRP-labeled goat anti-mouse IgG or IgG2c Fc antibody for 1hr (eBioscience). OD450 was measured on a microplate reader. Autoantibody activities against a panel of autoantigen specificities were measured using an Autoantigen Microarray platform developed by the University of Texas Southwestern Medical Center^69, 70^.

### Histology and Immunofluorescence Staining

Tissue specimens were fixed in 10% neutral buffered formalin and embedded in paraffin. Tissue sections were stained with periodic acid-Schiff (PAS) or with hematoxylin and eosin (H&E) and analyzed by light microscopy. The nephritis scoring system was adapted from the International Society of Nephrology/Renal Pathology Society (ISN/RPS) classification of human lupus nephritis. The final score accounted for morphological pattern (mesangial, capillary, membranous) and for the percentage of involved glomeruli. For immunofluorescence staining, kidneys or spleens were embedded in OCT and frozen in 2-methylbutane surrounded by dry ice. Frozen blocks were cut into 9μm section with cryotome and stored at -80°C. Upon thawing, sections were let dry at room temperature and stained. Spleen sections were stained with B220 (BD; RA3-6B2) and GL7 (BioLegend). Kidney sections were stained with FITC-labeled goat anti-mouse IgG (Jackson ImmunoResearch). Specimens were captured by Q capture software on a Nikon Eclipse microscope and quantifications were calculated using ImageJ software.

### RNA-seq Analysis

Quality of all RNA and library preparations were evaluated with BioAnalyzer 2100 (Agilent). Sequencing libraries were sequenced by the Epigenomics Core Facility at Weill Cornell Medicine using a HiSeq2500, 50-bp single-end reads at a depth of ∼15-50 million reads per sample. Read quality was assessed and adaptors trimmed using FASTP^71^. Reads were then mapped to the mouse genome (mm10) and reads in exons were counted against Gencode v27 with STAR2.6 Aligner^72^. Differential gene expression analysis was performed in R using edgeR3.24.3. Genes with low expression levels (<2 counts per million in at least one group) were filtered from all downstream analyses. Differential expression was estimated using quasi-likelihood framework.

Benhamini-Hochberg false discovery rate (FDR) procedure was used to correct for multiple testing. Genes with an unadjusted p-value less than 0.01 were considered differentially expressed. Downstream analyses were performed in R using a visualization platform built with Shiny developed by bioinformaticians at the David Z. Rosensweig Genomics Research Center at HSS.

Gene set enrichment analysis was performed using GSEA software (Broad Institute)^73^. Genes were ranked by the difference of log-transformed count per million (cpm) for contrasted conditions. Molecular Signatures DataBase v7.0 (Broad Institute) was used as a source of gene sets with defined functional relevance. Gene sets were also curated from RNA-seq datasets of ABCs from female DKO mice and from the blood of SLE patients^15, 20^ and from previously published PB/PC datasets^74, 75^. Gene sets ranging between 15 and 2500 genes were included into the analysis. Nominal p-values were FDR corrected and gene sets with FDR q <0.25 were used to crease GSEA enrichment plots. Analyses of differentially expressed genes were also performed using the online webtool CPDB and upstream regulator analyses were conducted using the Enrichr databases^76, 77, 78^.

### ATAC-seq Analysis

The nuclei of sorted ABCs from female DKO, male DKO, or Yaa-DKO mice were prepared by incubation of cells with nuclear preparation buffer (0.30M sucrose, 10mM Tris pH 7.5, 60mM KCl, 15mM NaCl, 5mM MgCl_2_, 0.1mM EGTA, 0.1% NP40, 0.15mM spermine, 0.5mM spermidine, and 2mM 6AA)^74^. Libraries were prepared as previously described^79^. Paired-end 50bp sequences were generated from samples with an Illumina HiSeq2500 and, following adapter trimming with FastP, were aligned against mouse genome (mm10) using bowtie2 with –local -q -p options. Peaks were called with MACS2 with *--macs2 callpeak -f BAMPE --nomodel --shit -100 --extsize 200 --B --SPMR -g $GENOMESIZE -q 0.01* options. Peak-associated genes were defined based on the closest genes to these genomic regions using RefSeq coordinates of genes. We used the *annotatePeaks* command from HOMER to calculate ATAC-seq tag densities from different experiments and to create heatmaps of tag densities. Sequencing data were visualized by preparing custom tracks for the UCSC Genome browser. *De novo* transcription factor motif analysis was performed with motif finder program *findMotifsGenome* from HOMER package on ATAC-seq peaks. Peak sequences were compared to random genomic fragments of the same size and normalized G+C content to identify motifs enriched in the targeted sequences.

### CUT&RUN Analysis

Freshly sorted ABCs and FoBs from both male and female DKO mice were used for the preparation of CUT&RUN libraries as described in ^80^ with slight modifications. We used three biological replicates for each condition. Sorted cells were immediately incubated overnight with H3K27me3 (anti-Tri-Methyl-Histone H3 (Lys27) (Cell Signaling Technology #Rabbit mAb #9733) or IgG isotype controls (Guinea Pig anti-Rabbit IgG (H+L) Secondary Antibody (novusbio #NBP1-72763) at 1:50 dilution of antibody binding buffer with a final concentration of 0.05% digitonin. Following incubation and digestion CUTANA™ pAG-MNase from EpiCypher (SKU: 15-1116) the digested DNA fragments were release and purified by QIAquick PCR purification kit (#2810). The resulting DNA fragments were end-repaired and barcoded libraries were made with the NEB Ultra II DNA Library Prep kit ((E7645S, E7335S & E7500S)). The libraries were sequenced on Illumina NextSeq 500 to a minimum of 15 million reads per sample. The data was analyzed using CUT&RUNTools pipeline ^81^ and resulting peak calls from SECAR were used for subsequent analysis. The peaks from IgG isotype controls were excluded in the SECAR peak calling program. A master file of H3K27me3 peaks across all the conditions and replicated was created and ncbi/BAMscale ^82^ was used to quantify peaks and generate scaled coverage tracks for viewing in UCSC genome browser. The raw counts on each H3K27me3 peak were obtained using ncbi/BAMscale cov –-bed master.H3K27me3.bed –prefix Peak_count –bam *.bam command and DESeq2 was employed to obtain differential H3K27me3 peaks between F-ABC and F-FOB mice. computeMatrix and plotHeap map functions were used from deepTools package to plot average tag densities and heatmap. Tag densities for the peaks in individual samples was calculated from DESeq2 and plotted as heatmap using Morpheus heatmap tool. The volcano plots were plotted with EnhancedVolcano^83^ available on Bioconductor and X chromosome heat of H3K27me3 signals was plotted using chromoMap. The z-statistic used in X chromosome H3K27me3 inactivation score heat map was obtained by implementing DESeq2 Wald test comparing F-ABC over F-FoB by taking log2 fold-change and dividing by its standard error resulting in z-statistic and a p-value is computed reporting that the probability that a z-statistic is small enough to reject null hypothesis. We summarized the differential peaks obtained by DESeq2 comparisions between F-ABC, F-FoB, M-ABC and M-FoB in Table S4.

### IgH Sequencing and Immune Repertoire Analysis

Genomic DNA was extracted from sorted cells using the QIAGEN Gentra DNA purification kit (Qiagen, No. 158689). Primer sequencies and library preparation were described previously^9^. Samples were amplified in duplicate (2 biological replicates per sample) using 100ng of input DNA per replicate (Table S4). Sequencing was performed on an Illumina MiSeq instrument in the Human Immunology Core facility at the University of Pennsylvania using a 2x300bp paired-end kit. Sequencing data analysis was also previously described^9^. Sequences were quality controlled with pRESTO^84^. Briefly, paired reads were aligned, sequences with low quality scores were discarded, and base calls with low confidence were masked with “N”s. IgBLAST was then used to align sequences to V- and J-genes in the IMGT database^85^. To group related sequences together into clones, ImmuneDB hierarchically clusters sequences with the same VH gene, same JH gene, same CDR3 length, and 85% identity at the amino acid level within the CDR3 sequence^86^. Clones with consensus CDR3 sequences within 2 nucleotides of each other were further collapsed to account for incorrect gene calls. Sequencing data were submitted to SRA under project number PRJNA663307 in accordance with the MiAIRR standard^87^. Prism v8.4.3 was used for D20, Jaccard index, SHM, CDR3 length, and VH gene usage histogram plots. Morpheus (Broad Institute) was used for VH heatmap. Venn Diagrams were generated in (http://www.interactivenn.net/). Other calculations were performed as described previously^86^.

### Statistics

*p-*values were calculated with two-tailed t-tests or ANOVA followed by multi-group comparisons, as indicated in the figure legends. Survival data was tested by Kaplan Meyer analysis with significance determined by the log-rank (Mantel-Cox) test. Correlation data was tested by paired Pearson correlations. *p-values* of <0.05 were considered significant. Ns: not significant, * p <0.05, ** p <0.01, *** p <0.001, **** p <0.0001. Statistical analysis was performed with Graphpad Prism 8.

## Data Availability

The data that support the findings of this study are available from the corresponding author upon request. The RNA-seq, CUT&RUN and ATAC-seq data will be deposited. All codes for bioinformatics analyses are available upon request.

## ACKNOWLEDGEMENTS

We thank members of the HSS Research Institute for thoughtful discussions and reagents. This work was supported by the US National Institutes of Health (AR064883 and AR070146 to ABP; T32 Rheumatology Research Training Grant to ER; P30 CA016520 and P30 AI0450080 to ELP), the Rheumatology Research Foundation, the Lupus Research Alliance, the Peter Jay Sharp Foundation, the Tow Foundation which provided support for the David Z. Rosensweig Genomics Research Center, Giammaria Giuliani and the Ambrose Monell Foundation, the Barbara Volcker Center for the Michael D. Lockshin Fellowship (MM), and Marina Kellen French and the Anna-Maria and Stephen Kellen Foundation (DJ and PS). Technical support was provided by the Epigenomics Core, the Microscopy and Imaging Core, and the Flow Cytometry Core Facility of Weill Cornell Medicine, by the Laboratory of Comparative Pathology at Memorial Sloan Kettering Cancer Center, by the Human Immunology Core at the Perelman School of Medicine, and by the Genomics and Microarray Core Facility at the University of Texas Southwestern Medical Center, and from the Office of the Director of the National Institutes of Health under Award Number S10OD019986 to Hospital for Special Surgery.

## SUPPLEMENTARY FIGURES

**Figure S1, related to Figure 1.**
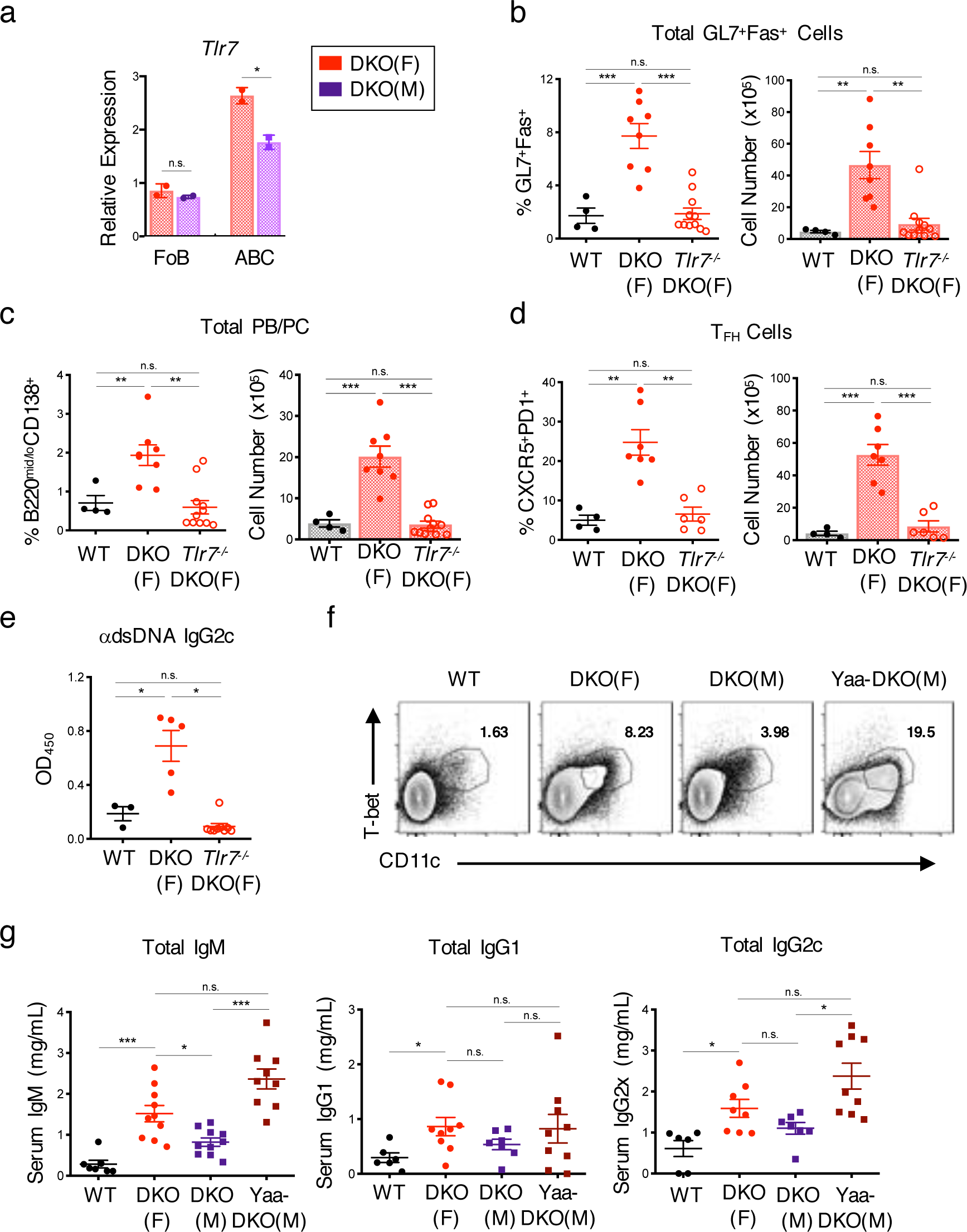
(A) Representative RT-qPCR of *Tlr7* expression in sorted ABCs (B220^+^CD19^+^CD11c^+^CD11b^+^) or follicular B cells (FoB; B220^+^CD19^+^CD11c^-^CD11b^-^CD23^+^) from female DKO (F) or male DKO (M) mice. Data representative of 3 independent experiments and show mean +/- SD; p-value by unpaired two-tailed t-tests. (B-D) Quantifications of total GL7^+^Fas^+^ B cells (*B;* B220^+^ GL7^+^Fas^+^), total PB/PC (*C;* B220^mid/lo^CD138^+^), and T_FH_ cells (*D;* CD4^+^ PD1^hi^CXCR5^+^) from the spleens of WT, DKO(F), and *Tlr7^-/-^.*DKO(F) mice. Data show mean +/- SEM; *n*>=4 per genotype; p-value by Brown-Forsythe and Welch ANOVA followed by Games-Howell’s test for multiple comparisons. (E) Pooled ELISA data for anti-dsDNA IgG2c autoantibodies in serum from the indicated mice. Data show mean +/- SEM; *n*>=3 per genotype; p-value by Brown-Forsythe and Welch ANOVA followed by Games-Howell’s test for multiple comparisons. (F) Representative FACS plots of CD11c^+^Tbet^+^ ABCs from the spleens of aged (20-30wk) WT, DKO(F), DKO(M), and Yaa-DKO(M) mice as in Fig. 1D. Data representative of at least 8 mice per genotype. (G) ELISA data for IgM, IgG1, and IgG2c antibodies in serum from the indicated mice. Data show mean +/- SEM; n*>=6* per genotype; p-value by Brown-Forsyth and Welch ANOVA followed by Games-Howell’s test for multiple comparisons. * p <0.05, ** p <0.01, *** p <0.001, **** p <0.001.

**Figure S2, related to Figure 2.**
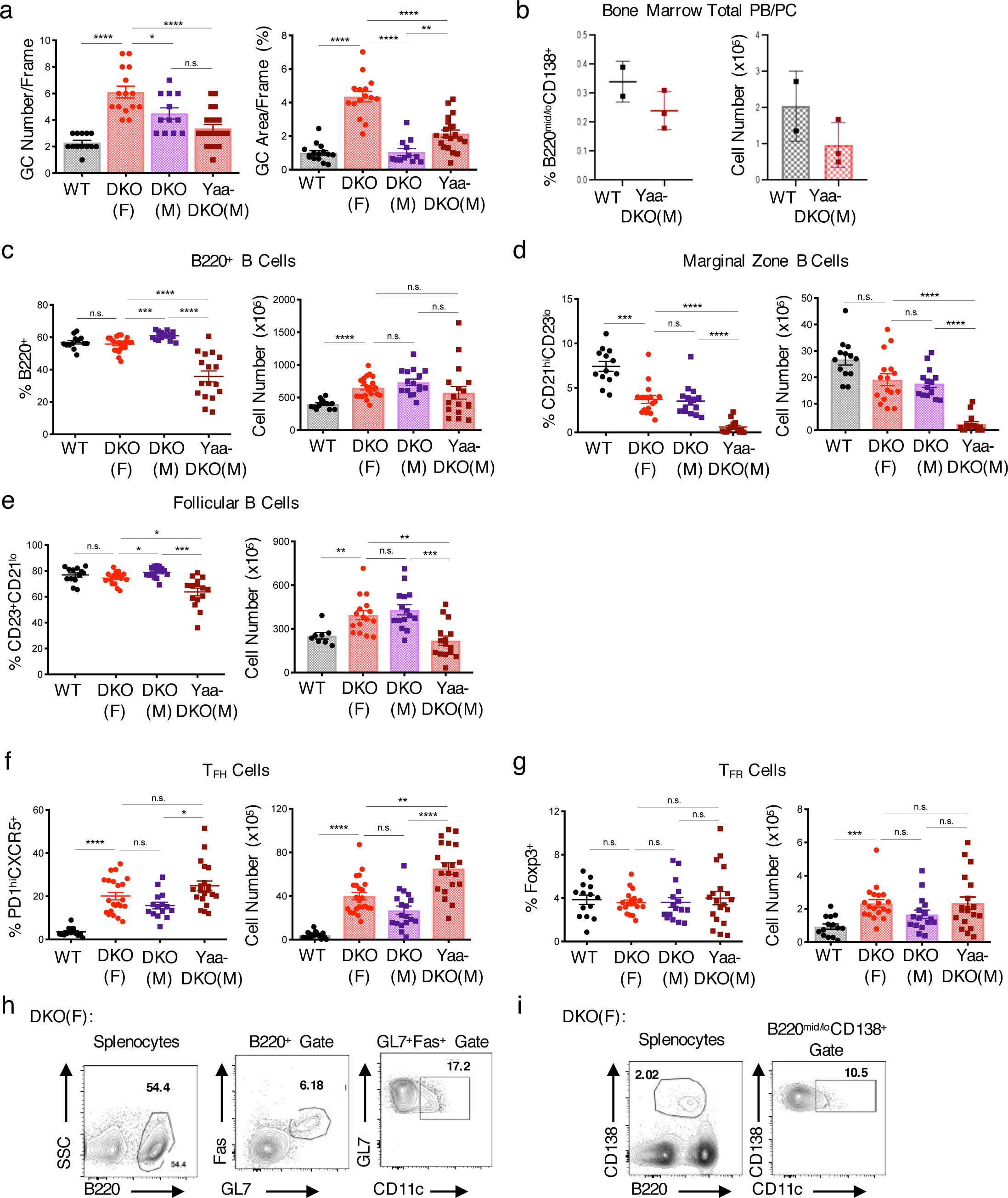
(A) Quantification of immunofluorescence images in Fig. 2B. Plots show GC count per frame *(left)* and GC area per frame (% GL7^+^B220^+^) *(right*). Data pooled from at least 4 frames for at least 2 mice per genotype and show mean +/- SEM; *n*>=10 per genotype; p-value by Brown-Forsythe and Welch ANOVA followed by Games-Howell’s test for multiple comparisons. (B-G) Quantifications of FACS data showing total bone marrow PB/PCs (B; B220^mid/lo^CD138^+^), total B cells (*C*; B220^+^ splenocytes), marginal zone B cells (*D*; B220^+^ CD21^hi^CD23^lo^ splenocytes), follicular B cells (*E*; B220^+^ CD23^+^CD21^lo^ splenocytes), T_FH_ cells (*F*; CD4^+^ PD1^hi^CXCR5^+^ splenocytes), and T_FR_ cells (*G*; CD4^+^ PD1^hi^CXCR5^+^ Foxp3^+^ splenocytes) from the indicated mice. Data show mean +/- SEM; *n*>=3 per genotype; p-value by Brown-Forsythe and Welch ANOVA followed by Games-Howell’s test for multiple comparisons. (H-I) Representative FACS plots showing the gating schematic for CD11c^+^ GL7^+^Fas^+^ cells in Fig. 2F *(H)* and for CD11c^+^ PB/PC in Fig. 2G *(I)* from DKO(F) mice. * p <0.05, ** p <0.01, *** p < 0.001, **** p <0.0001.

**Figure S3, related to Figure 3.**
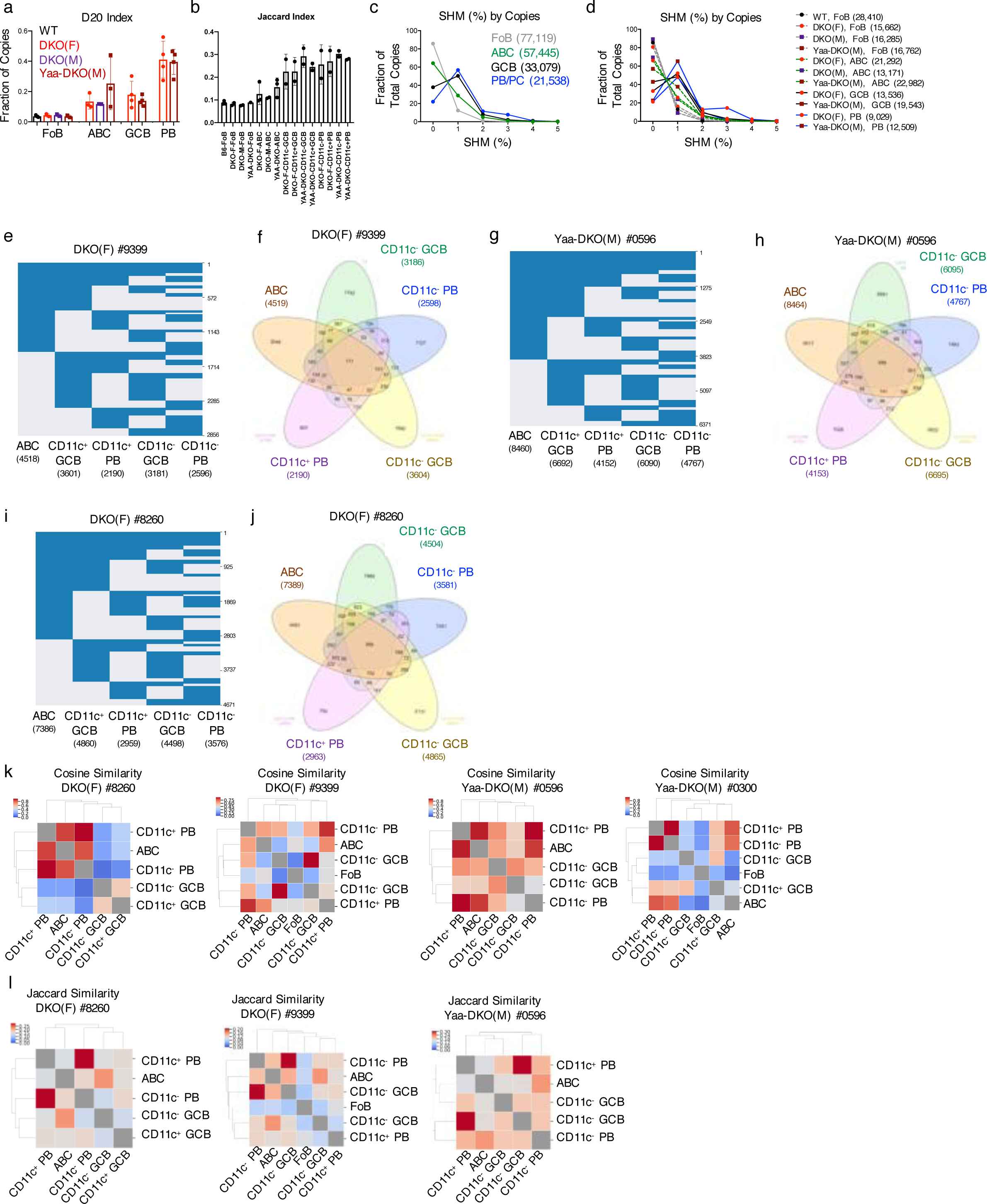
(A) D20 index of each mouse strain/subset combination. Each dot represents a mouse, the height of each bar shows the average across all mice, and error bars show SD. (B) Plot showing the level of overlap between clones in different strains/subsets. For each comparison, each clone was only counted once (no weighting for clone size) and functional overlap was computed using the Jaccard Index. Each dot represents a mouse/subset sample. (C-D) SHM data with sequence data weighted by copies (clone size). (E,G,I) Plot showing clones (rows) that overlap between at least two B cell subsets (columns). Numbers along the right side of the plot indicate clone counts. Data from the analysis of subsets from a single DKO female or Yaa-DKO male mouse as indicated. (F,H,J) Venn diagram showing clonal overlap where numbers indicate clone counts in the different subset interactions. Data from the analysis of subsets from a single DKO female or Yaa-DKO male mouse as indicated. (K) Heatmap showing Cosine similarity. Data from the analysis of a single DKO female or Yaa-DKO male mouse as indicated. (L) Heatmap showing Jaccard similarity (fraction of clones that overlap between different two-subset comparisons). Diagonal values are excluded for scaling. Data from the analysis of a single DKO female or Yaa-DKO male mouse as indicated.

**Figure S4, related to Figure 4.**
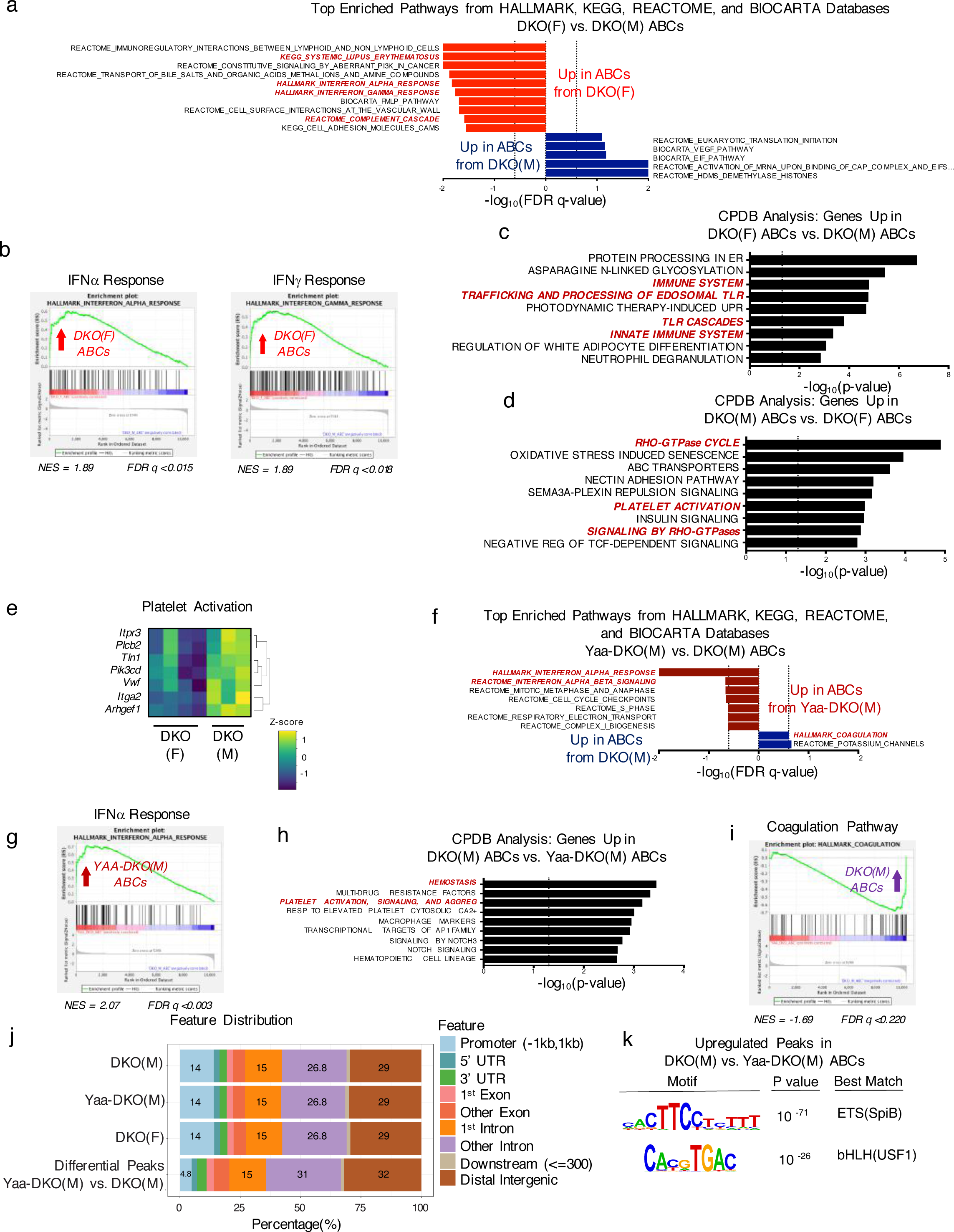
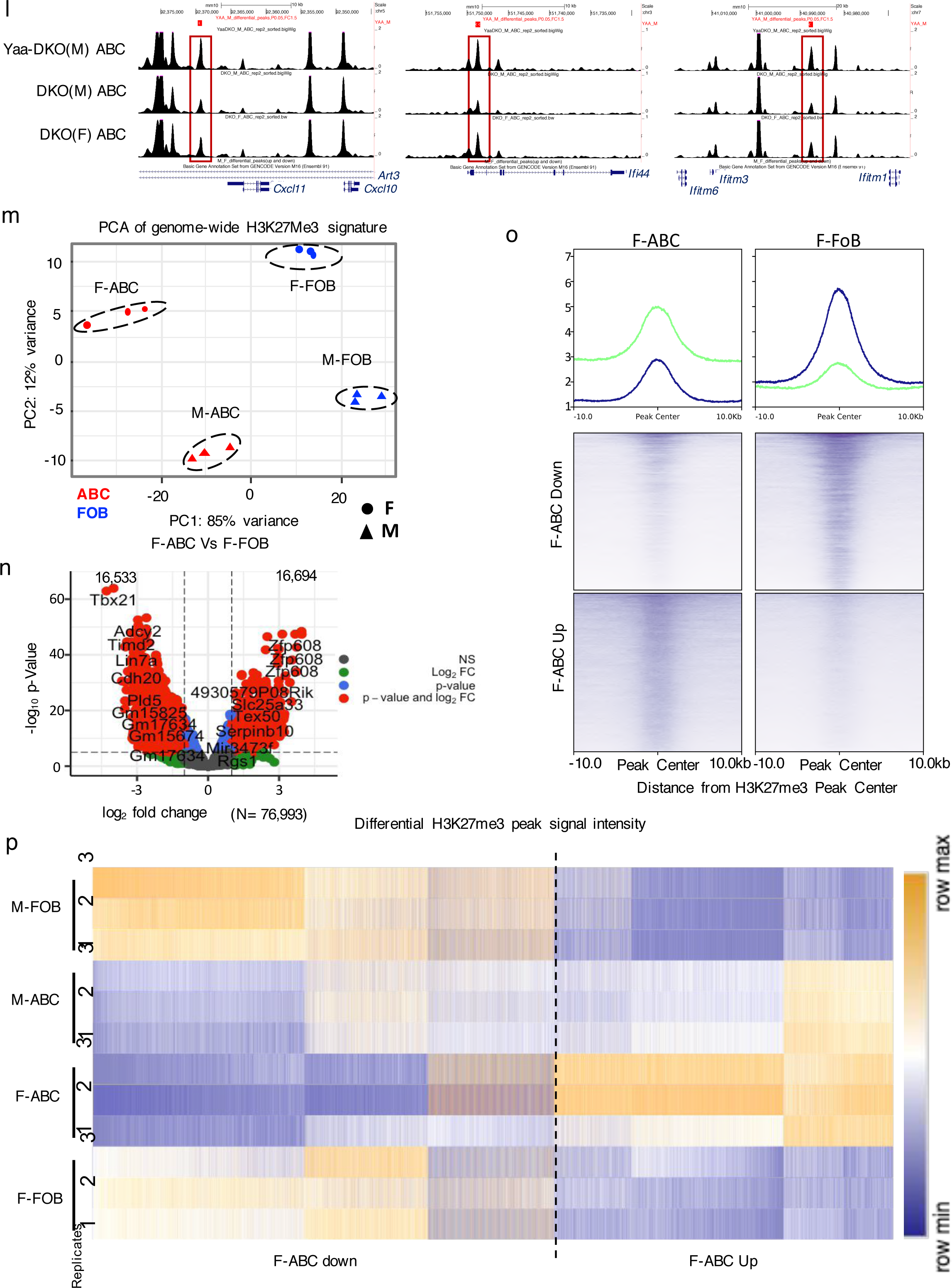
(A-I) RNA-seq was performed on sorted ABCs (CD19^+^B220^+^CD11c^+^CD11b^+^) from DKO(F), DKO(M), and Yaa-DKO(M) mice as in Fig. 4. (A) Plot showing the top pathways enriched in ABCs from DKO(F) *(red)* and DKO(M) *(blue)* mice as determined by GSEA using the HALLMARK, KEGG, REACTOME, and BIOCARTA databases. Dotted lines show significance thresholds at FDR q <0.25. (B) Plot showing enrichment of the HALLMARK_INTERFERON_ALPHA_RESPONSE and HALLMARK_INTERFERON_GAMMA_RESPONSE pathways in ABCs from DKO(F) mice. (C-D) Plots showing the top over- represented pathways comprising the genes significantly upregulated in ABCs from DKO(F) mice as compared to DKO(M) mice *(C)* or in ABCs from DKO(M) mice as compared to DKO(F) mice *(D)* using Consensus Pathway Database (CPDB). (E) Heatmap showing differentially expressed genes in the KEGG_PLATELET_ACTIVATION pathway. (F) Plot showing the top pathways enriched in ABCs from Yaa-DKO(M) *(maroon)* and DKO(M) *(blue)* mice as determined by GSEA using the HALLMARK, KEGG, REACTOME, and BIOCARTA databases. Dotted lines show significance thresholds at FDR q <0.25. (G) Table showing the top over-represented pathways comprising the genes significantly upregulated in ABCs from DKO(M) mice as compared to Yaa-DKO(M) mice using CPDB. (H) Plot showing the enrichment of the HALLMARK_INTERFERON_ALPHA_RESPONSE geneset in ABCs from Yaa-DKO(M) mice. (I) Plot showing the enrichment of the HALLMARK_COAGULATION geneset in DKO(M) ABCs as compared to Yaa-DKO(M) ABCs. (J) Plot showing the genomic distribution of peaks from ATACseq. (K) Motif enrichment analysis in ATAC-seq peaks that are significantly upregulated (logFC >1.5; p <0.05) in ABCs from DKO(M) mice. (L) Representative tracks showing accessible chromatin at the *Cxcl* cluster from DKO(F), DKO(M), and Yaa-DKO ABCs. (M) Principal component analysis of global H3K27me3 peaks. PC1 (85% variance) demarcates samples based on B cell phenotype and PC2 (12% variance) based on sex from CUT&RUN analysis. (N) Volcano plot showing differential H3K27me3 peaks between ABCs over FoBs from DKO(F) mice. A total of 76,993 peaks were identified of which 16,533/16,694 peaks were down/up- regulated by 2-fold and =< 0.05 p-value). (O) Average profile plot of H3K27me3 signal extended to -/+10 kb from the peak-center of 16,533 peaks in either downregulated in ABCs (*blue line*) or of 16,694 peaks upregulated in ABCs over FoBs from DKO(F) mice. Y-axis represents average tag density normalized to reads per million. A heatmap of the differential peaks of ABCs vs FoBs showing the spread of H3K27me3 signal from the peak center in ABC and FoB samples. (P) Heatmap depiction of tag densities of H3K27me3 peaks for down and upregulated peaks for individual replicates for ABCs and FoBs from male and female DKOs.

**Figure S5, related to Figure 5.**
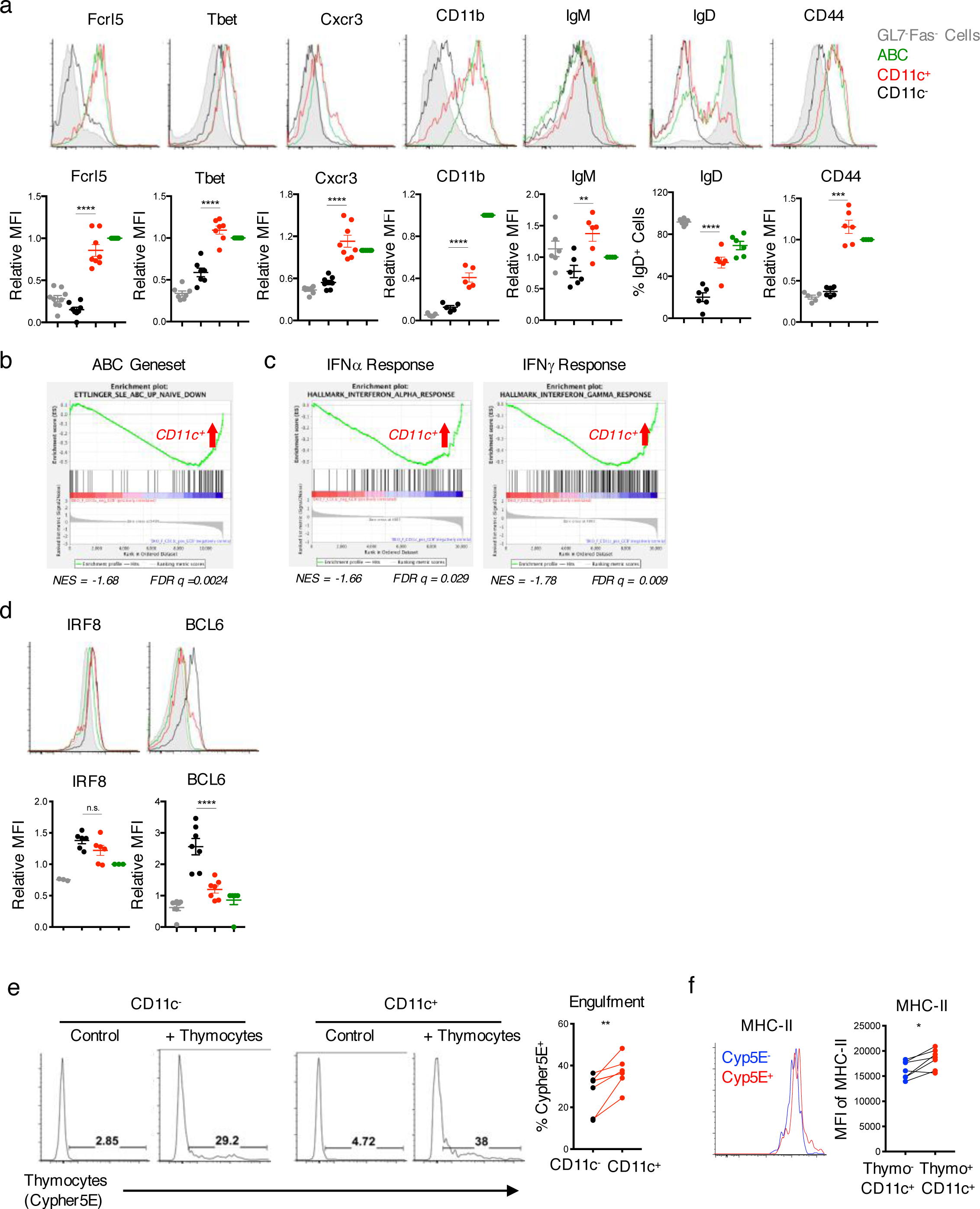
(A) Representative histograms and quantifications of Fcrl5, Tbet, Cxcr3, CD11b, IgM, IgD, and CD44 expression in CD11c^+^ CD19^+^GL7^+^Fas^+^ cells (CD11c^+^) (*red*), CD11c^-^ CD19^+^GL7^+^Fas^+^ cells (CD11c^-^) *(black)*, and CD19^+^CD11c^+^CD11b^+^ (ABCs) *(green)* from DKO(F) mice. CD19^+^Fas^-^GL7^-^ cells are shown as a control *(gray)*. Data representative of and/or pooled from at least 6 mice and show mean +/- SEM; p-value by 1-way ANOVA followed by Tukey’s test for multiple comparisons. (B) Plot showing the enrichment of an ABC geneset from SLE patients in CD11c^+^CD19^+^GL7^+^CD38^lo^ cells^15^. (C) Plot showing the enrichment of the HALLMARK_INTERFERON_ALPHA_RESPONSE and the HALLMARK_INTERFERON_GAMMA_RESPONSE genesets in CD11c^+^CD19^+^GL7^+^CD38^lo^ cells. (D) Representative histograms and quantifications of IRF8 and BCL6 in the indicated populations from DKO(F) mice. Data representative of and/or pooled from at least 6 mice and show mean +/- SEM; p-value by 1-way ANOVA followed by Tukey’s test for multiple comparisons. (E-F) Splenocytes from aged DKO mice were co-cultured for 3hr with Cypher5E-labeled thymocytes following induction of apoptosis with 50μM Dexamethasone. (E) Representative histograms and quantifications showing the percentage of CD11c^+^ and CD11c^-^CD19^+^GL7^+^Fas^+^ cells that engulfed apoptotic thymocytes (Cypher5E^+^). Data representative of and/or pooled from 4 DKO(F), 1 DKO(M), and 1 YAA-DKO(M) mice; p-value by paired two-tailed t-test. (F) Representative histogram and quantification of MHC-II expression in CD11c^+^CD19^+^GL7^+^Fas^+^ B cells that engulfed *(red*) or did not engulf *(blue)* apoptotic thymocytes. Data representative of and/or pooled from 6 mice as in Fig. 5E; p-value by paired two-tailed t-test.

**Figure S6, related to Figure 6.**
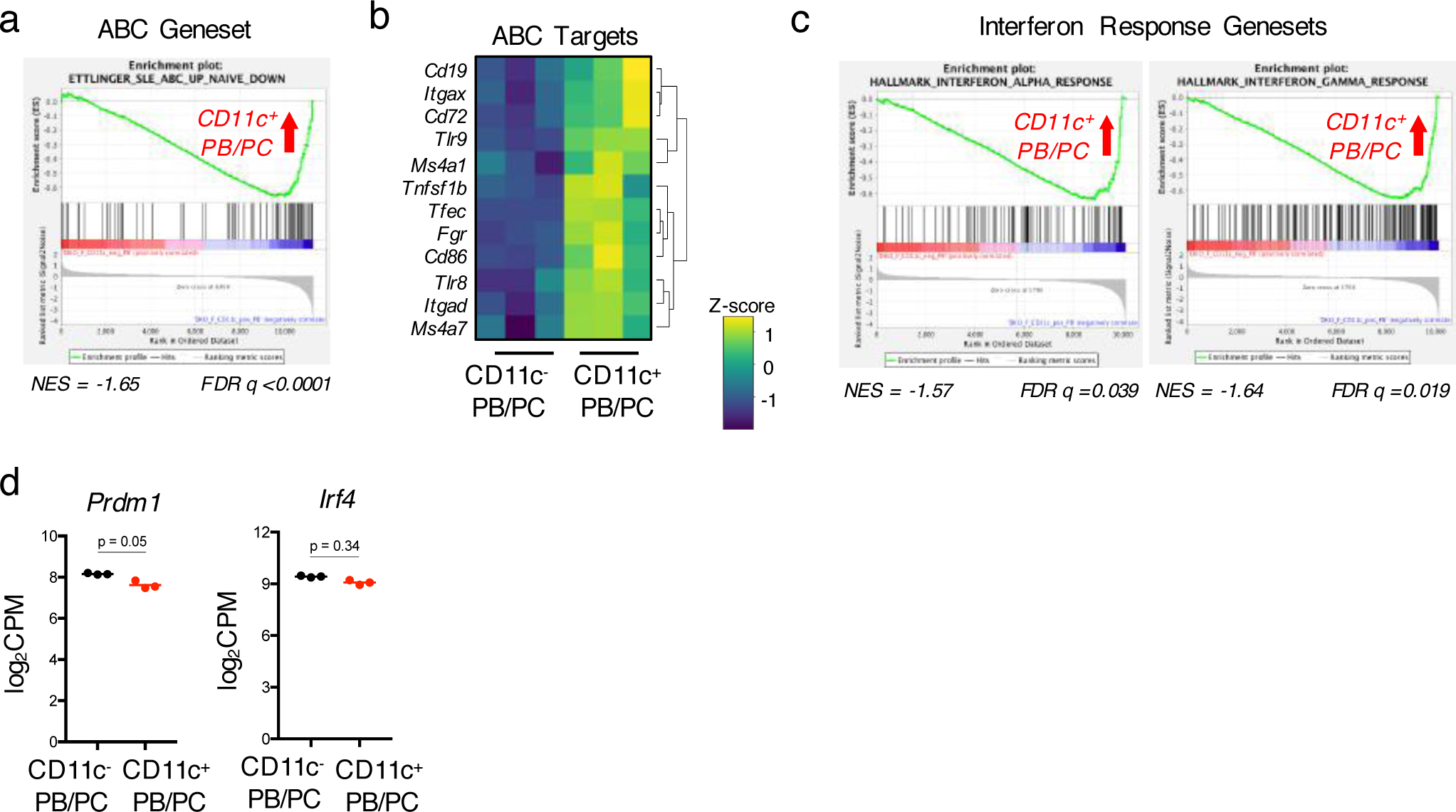
(A) Plot showing the enrichment of an ABC geneset from SLE patients in CD11c^+^ PB/PCs^15^. (B) Heatmap showing the differential expression of ABC target genes in CD11c^+^ and CD11c^-^ PB/PCs. (C) Plot showing the enrichment of the HALLMARK_ INTERFERON_ALPHA_RESPONSE and the HALLMARK_INTERFERON_GAMMA_RESPONSE genesets in CD11c^+^ PB/PCs. (D) Plots showing the normalized log2CPM counts for *Prdm1* and *Irf4* in CD11c^+^ and CD11c^-^ PB/PCs from Fig. 6A.

**Figure S7, related to Figure 8.**
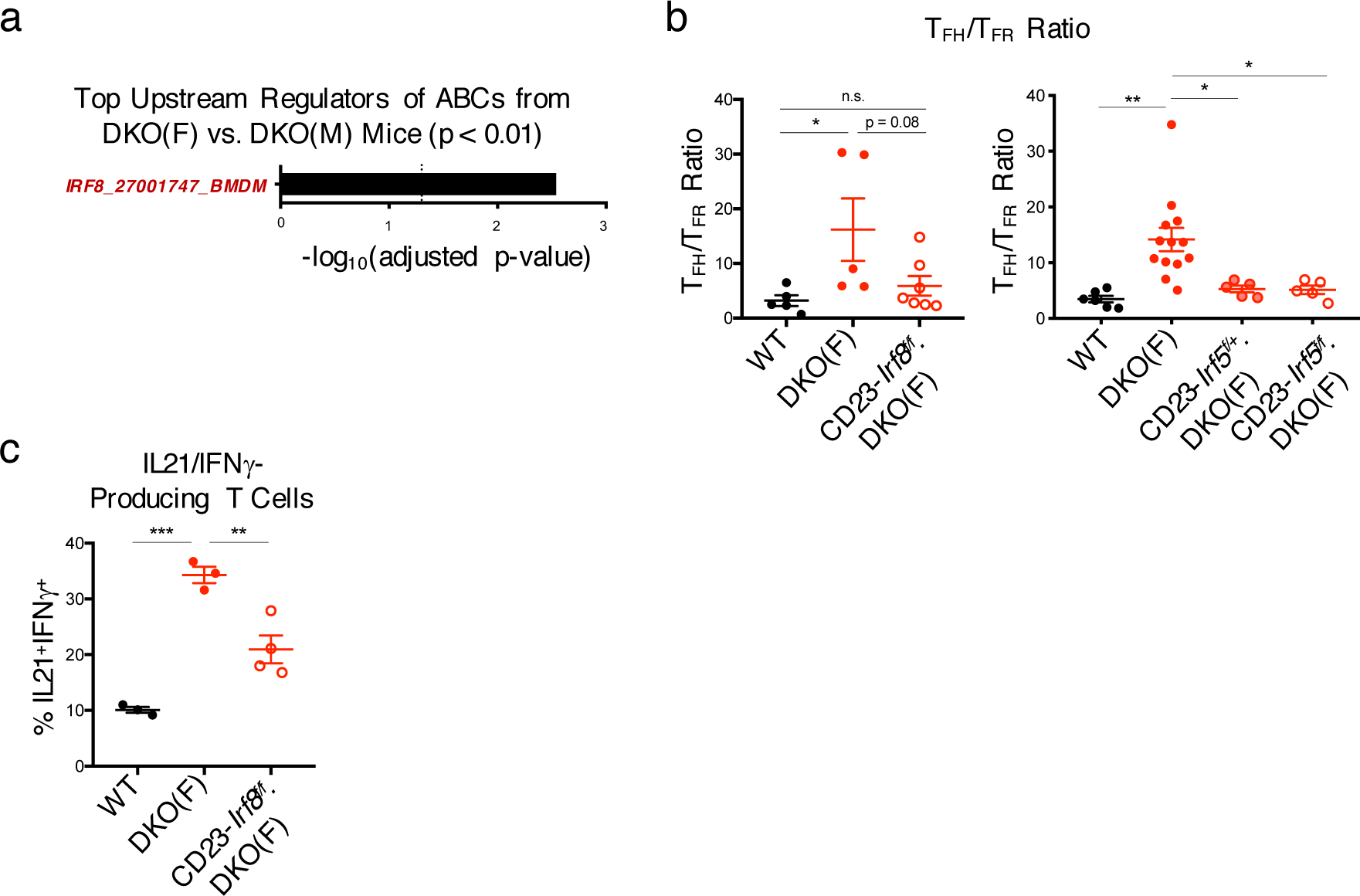
(A) Plot showing Enrichr analysis of potential upstream regulators for the geneset upregulated in ABCs from DKO(F) as compared to DKO(M) mice. (B) Quantifications showing the ratio of T_FH_ to T_FR_ cells from WT, DKO(F), CD23-Cre.*Irf8*^f/f^.DKO (CD23-*Irf8*^f/f^.DKO(F)), CD23-Cre.*Irf5*^f/+^.DKO (CD23-*Irf5*^f/+^.DKO(F)), and CD23-Cre.*Irf5*^f/f^.DKO (CD23-*Irf5*^f/f^.DKO(F)) mice. Data pooled from at least 5 mice per genotype and show mean +/- SEM; p-value by 1-way ANOVA followed by Tukey’s test for multiple comparisons. (C) Quantifications showing the production of IL21 and IFN*γ* in T cells from the indicated mice. Data pooled from at least 3 mice per genotype and show mean +/- SEM; p-value by 1-way ANOVA followed by Tukey’s test for multiple comparisons. * p <0.05, ** p <0.01, *** p <0.001, **** p <0.0001.

**Figure S8, related to Figure 9.**
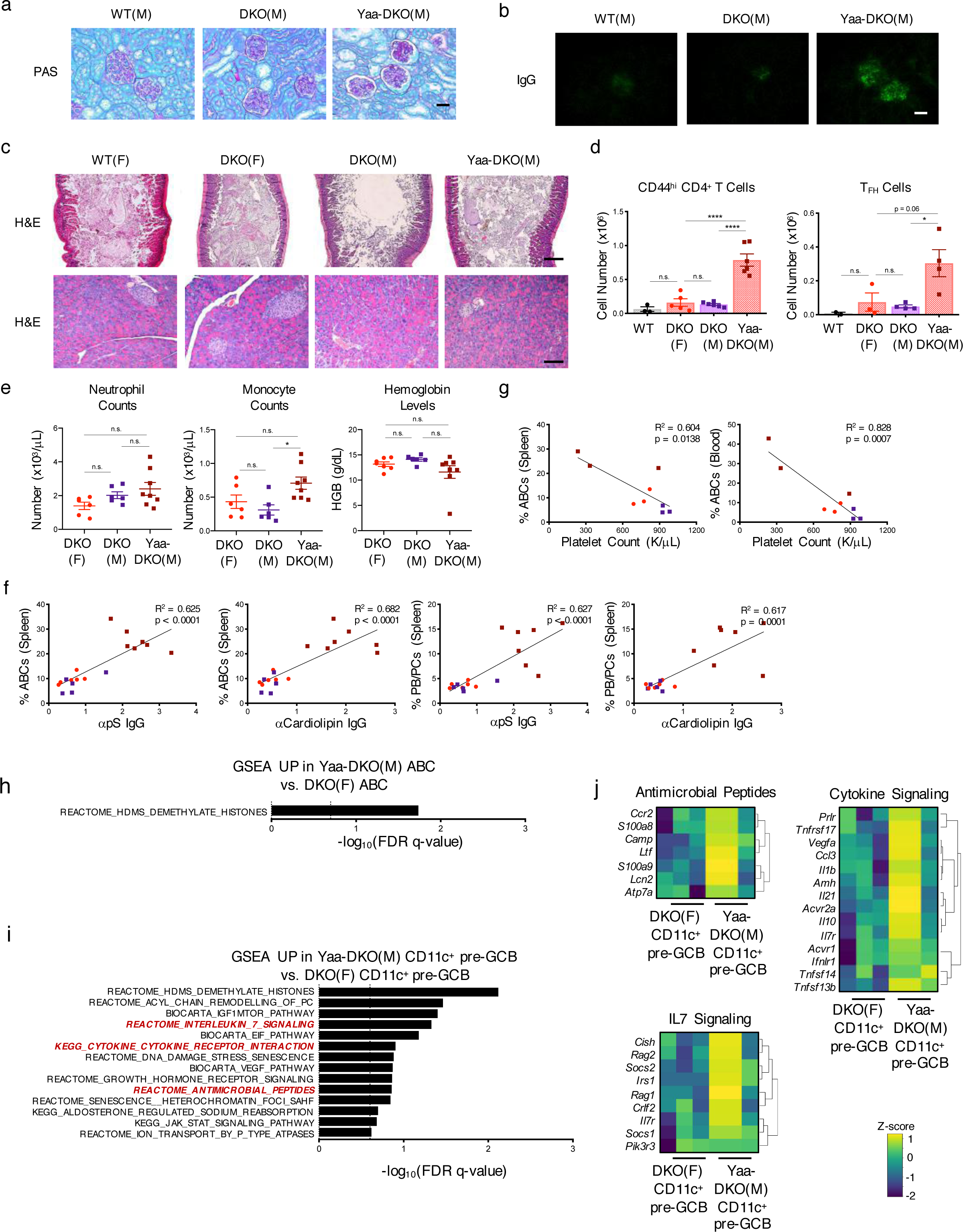
(A) Representative images of PAS staining in kidneys from the indicated mice. Data shows mean +/- SEM; *n>*=4 per genotype; p-value by 1-way ANOVA followed by Tukey’s test for multiple comparisons. (B) Representative images of IgG deposition in kidneys from the indicated mice. Data shows mean +/- SEM; *n*>=10 fields across 2-3 mice per genotype; p-value by 1-way ANOVA followed by Tukey’s test for multiple comparisons. (C) Representative H&E images of the colon *(top)* and pancreas *(bottom)* from the indicated mice. Data representative of 2 WT, 4 DKO(F), 4 DKO(M), and 4 Yaa-DKO(M) mice. (D) Quantifications showing the numbers of CD44^hi^ CD4^+^ T cells and T_FH_ cells (CD4+ PD1^hi^CXCR5^+^) in the lungs of the indicated mice. Data show mean +/- SEM; *n*>=2 per genotype; p-value by 1-way ANOVA followed by Tukey’s test for multiple comparisons. (E) Plots showing the numbers of neutrophils and monocytes and the levels of hemoglobin in the blood from the indicated mice. Data show mean +/- SEM; *n*>=6 per genotype; p-value by 1-way ANOVA followed by Tukey’s test for multiple comparisons. (F) Plots showing the correlations between the frequencies of ABCs or PB/PCs in the spleen and serum levels of anti-pS IgG and anti-Cardiolipin IgG antibodies. Data from DKO(F) *(red circles*), DKO(M) (*purple squares*), and Yaa-DKO(M) *(maroon squares*) are shown (*n>*=5 mice per genotype; p-value by Pearson correlation). (G) Plots showing the correlations between platelet counts and the frequencies of CD11c^+^CD11b^+^ ABCs in the spleen or blood. (H) Plots showing the top pathways enriched in ABCs from Yaa-DKO(M) mice as compared to those from DKO(F) mice as determined by GSEA. Dotted line indicates significance threshold at FDR q <0.25. (I-J) RNA-seq was performed on sorted CD11c^+^ GL7^+^CD38^lo^ (pre-GC) B cells from Yaa-DKO(M) mice. (I) Plots showing the top pathways enriched in CD11c^+^ pre-GC B cells from Yaa-DKO(M) mice as compared to those from DKO(F) mice as determined by GSEA. Dotted line indicates significance threshold at FDR q <0.25. (J) Heatmaps showing the expression of genes enriching the antimicrobial peptide, cytokine signaling, and IL7 signaling genesets in Yaa-DKO(M) CD11c^+^ pre-GC B cells. * p <0.05, ** p <0.01, *** p <0.001, **** p <0.0001.

